# Indecision and recency-weighted evidence integration in non-clinical and clinical settings

**DOI:** 10.1101/2024.11.11.622934

**Authors:** M del Río, N Trudel, G Prabhu, LT Hunt, M Moutoussis, RJ Dolan, TU Hauser

**Author notes:** Correspondence &. Shared senior authors (RJ Dolan, TU Hauser).

## Abstract

Biases in information gathering are common in the general population and can reach pathological extremes in paralysing indecisiveness, as in obsessive-compulsive disorder (OCD). Here, we adopt a new perspective on information gathering and demonstrate an information integration bias where most recent information is over-weighted by means of evidence strength updates (ΔES). In a large, crowd-sourced sample (N=5,237), we find reduced ΔES-weighting drives indecisiveness along an OC spectrum. We replicate the attenuated ΔES- weighting in a second lab-based study (N=105) that includes a transdiagnostic OC spectrum encompassing OCD and generalised anxiety patients. Using magnetoencephalography (MEG), we trace ΔES signals to a late neural signal peaking around 920 ms. Critically, high-OC participants show an attenuated neural ΔES signal in medio-frontal areas while other decision-relevant processes remain intact. Our findings establish biased information-weighting as a key driver of information gathering, where attenuated ΔES can lead to indecisiveness across an OC spectrum.

## Introduction

Knowing when to decide is difficult^1,2^. From selecting a movie on a streaming platform to buying a car, whenever there is uncertainty about the ‘right’ choice, gathering more information usually helps us make better decisions. But being indecisive and spending too much time gathering information is also disadvantageous and carries substantial opportunity and energy costs^3^.

In humans information gathering biases are common, often expressed as gathering too little or too much information^4,5^. Dramatically inflated information gathering biases are a feature of psychopathology and believed to drive core symptoms^4,6–11^. For example, indecisiveness and persistent doubt can be debilitating aspects of obsessive-compulsive disorder (OCD)^12,13^. Excessive deliberation also extends beyond core OCD symptoms and can impact everyday wellbeing and function^14^.

While indecisiveness in OCD is traditionally assessed using clinical interviews^15^, recent literature has provided a more quantitative approach based upon using information gathering tasks^4,10,16–23^. Studies using the latter have shown that patients with OCD, as well as nonclinical participants with high obsessive-compulsive (OC) scores, manifest elevated levels of information gathering before committing to a decision. While similar effects have been observed in naturalistic settings^23^, the precise neurocognitive mechanisms underlying the expression of indecisiveness remain unknown.

A relevant field of study, largely disconnected from information gathering, is that of evidence accumulation in decision making^24^. Here it is considered that evidence for different options increases gradually with incoming information, and a decision is made when evidence for an option hits a decision threshold^24–29^. The latter is believed to collapse with time, meaning that decisions become increasingly liberal, which is often framed in terms of an urgency signal^30–33^. These effects are observed in a range of information gathering contexts despite striking differences in tasks and timescales^34–36^. Importantly, recent studies suggest that evidence does not accumulate linearly as a function of decision-relevant information, but in a recency-biased manner such that most recent evidence is over-weighted^37–40^. Determining whether such biases also exist in information gathering, and whether they act as drivers for indecisiveness is an important question.

In this study, we investigate the neurocognitive contributors to information gathering across two large samples and tasks which we previously collected in the lab. We find that, in addition to urgency-like signals, evidence is integrated non-monotonically, with what we term evidence strength updates (ΔES) critically determining when a participant decides to decide. Furthermore, across both studies, we find that participants along an OC spectrum rely less on these ΔES both in clinical and nonclinical samples. Using magnetencephalography (MEG), we show this attenuation of ΔES is mirrored in the brain, with neural ΔES representations in medio- frontal areas being less evident in OC whereas other critical contributors to information gathering remain intact. We conclude that indecisiveness along an OC spectrum is driven by a reduced reliance on most recent information.

## Results

### Cognitive contributors to information gathering

To first ascertain a link between information gathering and OC symptoms in the general population, we conducted a large-scale, crowd-sourced study (N = 5,237 at the time of analysis). To this end, we implemented an information gathering task on an app for handheld electronic devices (www.brainexplorer.net), where participants had to decide which of two stimuli was more plentiful across 25 hidden locations (Fig. 1A). Participants were free to sample as many of the locations as they wanted before committing to a decision, such that we could use the number of draws prior to a decision as an index of indecisiveness.

**Figure 1:**
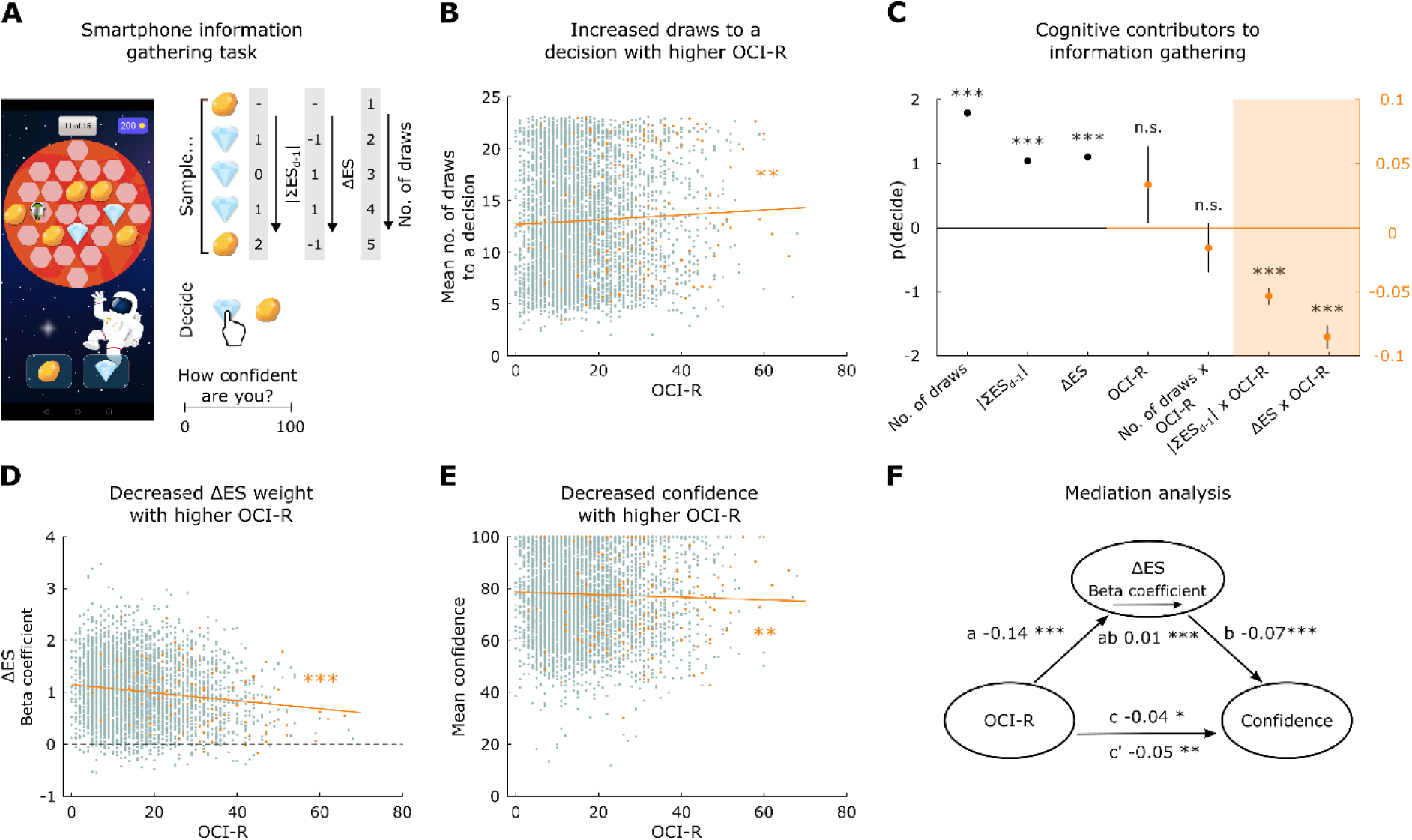
OC symptoms are linked to increased information gathering in a large-scale smartphone population study. **A** The participants’ task was to determine which of two possible gems was more abundant. They were free to uncover as much evidence as they wished, by tapping on locations on the grid, until they decided to commit to a binary choice between one of two gems. An example sequence of samples and corresponding evidence accumulation process for cumulative evidence strength at the previous draw |ΣESd-1| and the evidence strength update ΔES is depicted on the right. Participants rated their confidence after every binary choice, then received 100 points for correct responses or lost 100 points for incorrect responses. **B** Within a large-scale sample of the general population, individuals with higher OC symptoms sampled more information before committing to a decision. Data points pertaining to individuals with a self-reported OCD diagnosis are shown in orange. **C** The probability of making a decision p(decide) was predicted from experimental factors as well as individual differences in OC symptoms using a general linear mixed model. The experimental factors (left y-axis) comprised current draw number, cumulative evidence from the start of the game until the previous draw (|ΣESd-1|), and the evidence strength update (ΔES). More draws and higher |ΣESd-1| and ΔES increased the probability of making a decision p(decide). Individual differences in obsessive-compulsive symptoms (right y-axis) were quantified as the main effect of the OCI-R score, and the interactions of the OCI-R score with the experimental factors. The analysis showed that OC symptoms per se did not have a significant effect on p(decide), and that people with high OC symptoms did not weight the current draw number differently in deciding to commit to a choice – instead they weighted both previous and current evidence less. The latter effect is illustrated in **D**: to obtain estimates of sensitivity to evidence at a single-participant level, we fit a GLM predicting p(decide) separately for each participant. Convergent with the significant interaction effect in the GLMM, those with higher OC symptoms had reduced beta coefficients for ΔES, i.e., they weighed ΔES less in deciding to commit (N = 3,903 after removing outliers, defined as any participant with any beta values outside of the interquartile range). **E** Individuals with higher OC symptoms also rated their confidence in their decisions as being lower on average (N = 3,293, see exclusion criteria in the Methods). **F** A mediation analysis suggests that sensitivity to ΔES partially mediates the association between OC symptoms (OCI-R total score) and confidence ratings, such that lower confidence is partially explained by altered evidence integration. Asterisks denote the level of significance (* p < 0.05, ** p < 0.01, *** p < 0.001) and error bars in **C** the standard error of the mean (fixed effects error bars hidden by the mean itself).

First, to better understand the cognitive contributors to participants’ information gathering behaviour, we used general linear mixed models (GLMMs) to predict on each draw whether a participant would continue sampling information or make a decision (*p(decide)*). In this task, information accumulated non-monotonically, allowing us to distinguish the contribution of different factors (e.g., current information, total evidence, number of draws) to commitment to a decision. To test whether a recency bias^41–43^ is present in an information gathering context, we distinguished total evidence strength on the previous draw (|ΣESd-1|; absolute cumulative evidence difference across draw 1 to d-1 for the current majority) from the most recent change in evidence strength or evidence strength update (ΔES; Fig. 1A).

Model comparison favoured the aforementioned distinction over a model where all evidence was aggregated in a single predictor (ΔAIC = 1.709 × 10^6^; ΔBIC = 1.709 × 10^6^). We found that both the current update in signal strength (ΔES, β = 1.098, SE = 0.008, p < 0.001; Fig. 1C) and previous evidence strength (|ΣESd-1|; β = 1.038, SE = 0.005, p < 0.001) positively predicted whether participants would commit to a decision. An additional GLMM predicting *p(decide)* from the current evidence at draw d, d-1 and d-2 relative to the chosen option showed a decreasing contribution of current evidence for lagged draws (ESd: β = 0.648 SE = 0.007, p < 0.001; ESd-1: β = 0.253 SE = 0.004, p < 0.001; ESd-2: β = 0.195 SE = 0.005, p < 0.001, see Supplementary Fig. 1A). This accords with a recency-bias in information gathering, analogous to that found across other decision making domains^37,38,57,58^. It is particularly striking as this task was designed so that all information remained present on the screen irrespective of when it was gathered, such that this recency-bias is not accounted for by working memory limitations or related explanations.

In keeping with the idea of collapsing decision boundaries or urgency signals^4,30,44,45^, we added draw number as a predictor. Draw number additionally predicted whether participants would make a decision (β = 1.799, SE = 0.006, p < 0.001) such that participants were more likely to stop gathering information the longer they had already spent gathering information (irrespective of how strong the evidence is) and, in addition, if they (recently) received stronger evidence favouring one option over the other.

### OC-linked indecisiveness is due to attenuation of evidence weighting

Next, we investigated whether an OC spectrum was linked to indecisiveness. Taking the number of draws prior to making a decision as an indicator of indecisiveness^4,34,36,46,47^, we found - as hypothesised - that participants with higher OC symptoms (as determined by the OCI-R total score^48^) gathered more evidence before making a decision (Fig. 1B, ρs = 0.040, p = 0.004), while not showing any difference in accuracy for these decisions (ρs = -9.998 × 10^-4^, p = 0. 942). Thus, in this large information-gathering dataset we find indecisiveness relates to OC symptoms across the general population.

Next, we investigated which of the identified cognitive contributors explained differences in task behaviour in high-OC participants. We found that OC symptoms were linked to reduced weighting of previous evidence |ΣESd-1| (β = −0.053, SE = 0.007, p < 0.001) and particularly of ΔES (β = -0.085, SE = 0.009, p < 0.001). There was no main effect of OC symptoms on sampling decisions (*p(decide)*; Fig. 1D, β = 0.033, SE = 0.030, p = 0.265), implying that the increased draws prior to a decision (Fig. 2B) is not a simple consequence of a tendency not to decide; and, contrary to our initial assumptions^4,49^ there was no effect on urgency (as captured by the interaction with the number of draws; β = -0.016, SE = 0.019, p = 0.265). We found similar but weaker effects of OC symptoms on evidence weighting when comparing a subset of participants with a self-reported OCD diagnosis with those who explicitly declared no lifetime psychiatric disorder (see Supplementary Materials). Further, by fitting an additional GLM predicting *p(decide)* from the current evidence at draw d, d-1 and d- 2 relative to the chosen option per participant, we found that a difference in beta weights for the evidence at draw d and the evidence at draw d-1 correlated negatively with the OCI-R score (ρs = -0.068, p < 0.001), supporting the notion of a decreased recency effect. This means that high-OC participants were no more likely to decide per se, or express an altered urgency signal, but instead showed an attenuated impact of the most recent information sample. In other words, evidence strength updates seem to be driving an indecisiveness along the OC spectrum.

**Figure 2:**
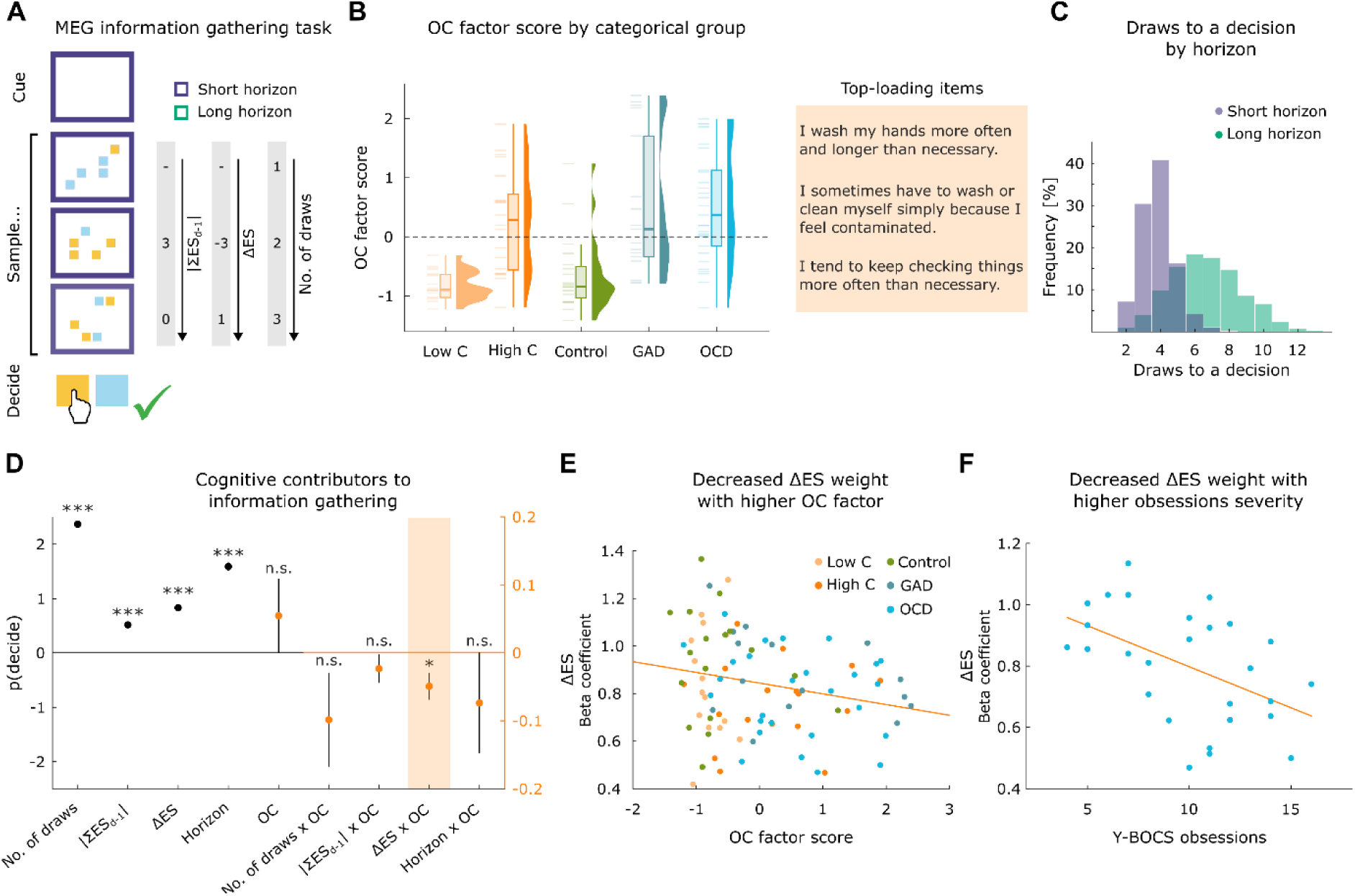
Reduced ΔES weighting replicated in transdiagnostic dimensional patient-control sample. **A** Participants’ goal was to determine which of two possible stimuli was more abundant. They were presented with a sequence of draws until they decided to declare for a stimulus. Each draw consisted of five stimuli at a time in varying proportions. Here, sequences were constructed to reduce the collinearity between the number of draws, current and previous evidence, which will allow us to study their specific representations in the brain. The maximum number of draws varied between 4 and 8 in the short horizon condition and between 10 and 14 in the long horizon condition. The horizon condition was cued by the colour of the frame (different colours were presented in the task). Participants received 2 points for a correct response, lost 2 points for an incorrect response, and lost 1 point for non-decisions, which were later translated into bonus payments. **B** In order to pool across clinical and non-clinical groups, we reduced the dimensionality of seven questionnaires using an exploratory factor analysis. This revealed a three-factor solution, where the second factor mapped primarily onto items in questionnaires assessing OCD (OCI-R and PI-WSUR). We used this OC factor to quantify individual differences across all participants. **C** The number of draws to a decision varied across trials and participants in the short and long horizon condition. Note that the number of draws to a decision per se was not associated with the transdiagnostic OC factor (ρs = -0.117, p = 0. 236), yet this is not surprising given the highly constraining horizon manipulation. **D** The probability to make a decision was predicted in a GLMM from experimental factors as well as variability in the OC factor. Experimental factors (left) comprised the number of draws, the cumulative evidence strength from the start of the game until the previous draw|ΣESd-1|, ΔES, and the horizon condition (controlling for interactions of the horizon condition with all other experimental factors). More draws, higher previous and current evidence strength, and a short horizon condition all increase the probability to make a decision. Individual differences in the OC factor (right) are quantified as the main effect of the OC factor score and its interaction with the experimental factors. The GLMM shows no significant effect of the OC factor per se, or differences in weighting the current number of draws or the previous evidence in making a decision. Conversely, it shows that those with higher OC factor scores weight ΔES less in making a decision. **E** Illustration of the association between ΔES weighting and the OC factor (using individual GLMs per participant; N = 92 after outlier exclusions were applied to all beta coefficients in the GLM). **F** OCD patients with stronger obsession symptom severity had more attenuated weighting of ΔES (ρs = -0.475, p = 0.012). Asterisks denote the level of significance (* p < 0.05, ** p < 0.01, *** p < 0.001).

### Lowered confidence in high-OC participants mediated by attenuated ΔES

Having identified key drivers of indecisiveness, we next assessed if this was a purely behavioural effect or whether it also affected participants’ perception of their performance or insight. To assess this, we asked participants to rate their post-choice confidence (prior to viewing the outcome; Fig. 1A). We found that OC symptoms were also linked to confidence, with high-OC participants having lower confidence (Fig. 1E; ρs = -0.054, p = 0.002, N = 3,293 after excluding additional participants based on confidence data criteria, see Methods). This means that the OC spectrum was not only linked to indecisive information gathering but also to a lack of confidence, a finding that aligns with evidence of impoverished confidence in other domains^50^. It also chimes with the notion that indecisiveness is closely linked to subjective doubt.

Lastly, using a mediation analysis^51,52^, we evaluated the link between the reduced ΔES weight from our information gathering analysis and confidence findings. We found that the ΔES weighting (using beta estimates from non-hierarchical models, Fig. 1D) was associated with task confidence (b = -0.07, SE = 0.018, t(3293) = -3.754, p < 0.001, Fig. 1F). More specifically, we found that ΔES partially mediated the influence of OC symptoms on confidence (ab = 0.01, SE = 0.003, t(3293) = 3.402, p < 0.001, c = -0.04, SE = 0.017, t(3293) = -2.467, p = 0.014, c’ = -0.05, SE = 0.018, t(3293) = -2.986, p = 0.003, Fig. 1F). This means that the lower confidence in high-OC participants is, at least in part, influenced by the same underlying process as that of attenuated ΔES.

### Cognitive contributors to information gathering replicate in a lab-based sample

As the above dataset was primarily derived from a large, self-selected, population of non-clinical participants, we next asked whether the findings also held true in a more targeted patient sample. Thus, we analysed an independent sample that included patients assessed in the lab (N = 105), using a novel adaptation of the above information gathering paradigm (Fig. 2A) which had been previously collected in the lab.

We first replicated the findings related to the cognitive contributors to information gathering per se (Fig. 2C). Both ΔES (β = 0.830, SE =0.020, p < 0.001) and previous evidence strength (β = 0.516, SE = 0.021, p < 0.001) positively predicted when participants would make a decision. Likewise, we replicated the effect of current draw number (β = 2.367, SE = 0.069, p < 0.001), supporting the notion of an emerging decision urgency. A noteworthy extension here was that this lab version featured two different finite horizons, meaning that each game would end after a probabilistically predetermined number of draws (either 4-8 in the ‘short’ or 10-14 in the ‘long’ horizon condition). We found that this horizon condition significantly predicted participants’ decision making (β = 1.602, SE = 0.074, p < 0.001). In other words, participants are not only more likely to commit to a decision if they (recently) received more evidence for one of the options and if they had already gathered information for longer, but also if they are incentivised to trade off speed for accuracy.

### Attenuated ΔES are also linked to OC symptoms in a transdiagnostic clinical sample

Next, we investigated how these predictors’ link to an OC spectrum in a sample consisting of a clinical group of OCD patients (N = 29) and generalised anxiety disorder (GAD) patients (N = 17), as well as non-patient cohorts with varying degrees of OC symptoms (high OC: N = 20, low OC: N = 20, controls: N = 19). This recruitment strategy allowed for a broad sampling of OC symptoms not only in OCD patients but also in adjacent disorders as well as non-clinical high-OC participants. A similar approach has previously revealed striking dissociations between transdiagnostic dimensions and traditional diagnostic categories^53^. To derive a transdiagnostic and dimensional OC symptom measure, we factor analysed key psychiatric questionnaires collected across the entire sample, yielding a three-factor solution where one factor primarily reflected OCD and related symptoms (see Methods and Supplementary Fig. 2). On average, OCD patients ranked highest on this factor, followed by high-compulsive non-patients and GAD patients, while low-compulsive and controls ranked the lowest (see Fig. 2B).

Investigating how this transdiagnostic OC factor linked to cognitive contributors of information gathering, we again replicated the effect of ΔES (β = -0.049, SE = 0.020, p = 0.013, Fig. 2C and D), such that high-OC participants weighted the update in evidence strength less in their decision to declare for a stimulus. Similar, albeit less robust, effects were seen when comparing OCD patients to matched controls alone (β = -0.102, SE = 0.061, p = 0.046, one-sided test). In addition, by fitting an additional GLM predicting *p(decide)* from the evidence at draw d, d-1 and d-2 relative to the chosen option per participant, we again found a decreasing contribution of current evidence for lagged draws (ESd: β = 0.519 SE = 0.015, p < 0.001; ESd- 1: β = 0.212 SE = 0.013, p < 0.001; ESd-2: β = 0.216 SE = 0.010, p < 0.001, see Supplementary Fig. 1B), and that the difference in beta weights for the evidence at draw d and the evidence at draw d-1 correlated negatively with OC symptoms (ρs = -0.199, p = 0. 024, one-sided test).

As was the case for the previous population sample, we find no main effect of OC symptoms (β = 0.055, SE = 0.055, p = 0.318), or any interaction with the current number of draws (β = -0.098, SE = 0.069, p = 0.155) or with the horizon condition (β = -0.074, SE = 0.074, p = 0.320). In this sample, the interaction of OC symptoms with previous evidence did not reach significance (β = -0.023, SE = 0.021, p = 0.272), in contrast to the first, larger sample. In sum, this further supports the notion that an over-reliance on most recent information during information gathering is attenuated in high-OC participants, including patients.

### Attenuated ΔES are primarily linked to obsessions in OCD patients

Lastly, we asked whether an attenuated weighting of ΔES was linked to symptom severity, especially in patients with OCD. To this end, we correlated the scores from the clinical Y-BOCS interview (only conducted in OCD patients by trained researchers) with the ΔES effect on information gathering. Interestingly, we found that attenuated weighting of ΔES was significantly associated with symptom severity for obsessions (ρs =-0.475, p=0.012), but not for compulsions (ρs = -0.205, p = 0.306; total score trended at ρs = -0.342, p = 0. 081). This suggests that the attenuated weighting of recent information may be primarily linked to an obsession dimension, in line with the idea that underweighting of information about the unlikely nature of the obsessions promotes their perpetuation.

To assess how generalisable this effect was, we went back to the complete samples in both studies, and found that ΔES attenuation related to obsessions from the OCI-R questionnaire both in the smartphone population sample (ρs = -0.083, p < 0.001) as well as in the lab-based sample (including patients and non-patients: ρs = -0.226, p = 0.030). However, these effects were less specific (see Supplementary Fig. 3), possibly due to range effects across the wide spectrum of patients and non-patients.

Thus, our findings indicate that attenuated ΔES are not only associated with OC symptoms across a wide OC spectrum, but also linked to symptom severity of obsessions within OCD patients.

### The brain processes decision-relevant factors sequentially

To better understand the brain processes relevant for information gathering, we concurrently acquired magnetoencephalography (MEG) data from all participants in our second study. Here the validity of our behavioural model leads to the prediction that we could decode (5-fold cross-validated iterative lasso regression with whole-brain MEG^54^, see Methods) each of the relevant cognitive factors used to predict information gathering, as described above - namely, ΔES, total evidence strength at draw d-1 |ΣESd-1|, horizon, and number of draws.

We found evidence for a representation of each of the aforementioned factors in our MEG data (Fig. 3A; p < 0.05 cluster-based permutation tests at t > 3), supporting the idea that the brain tracks these variables. Interestingly, these representations were instantiated at different times within a trial following a sequential order, where pre-existing latent factors were represented first and novel information, that needed to be computed based on these latent representations, was represented later in a trial. Specifically, we found that the representations of the number of draws, horizon and the total evidence at draw d-1 preceded the representation of ΔES (comparison of maximum decodability timepoint; number of draws vs ΔES: t(104) = - 14.773, p < 0.001; horizon vs ΔES: t(104) = -5.387, p < 0.001; |ΣESd-1| vs ΔES: t(104) = -23.750, p < 0.001, one-sided tests; Fig. 3B). Importantly, the representation of the currently shown evidence (quantified as the relative evidence for the two alternative response options in the current draw) also preceded the representation of ΔES (t(104) = -8.711, p < 0.001, see Supplementary Fig. 4C), as would be expected when considering that ΔES is computed based on both previous total evidence and current evidence. In addition, we observed that latent variables that are not dependent on currently presented information manifest as a more sustained representation (e.g., horizon), consistent with the fact they do not need to be computed on the fly after each draw. Thus, our findings demonstrate a cascade of neural information processing and representation that aligns with the evolution of cognitive processes we deciphered from participants’ behaviour.

**Figure 3:**
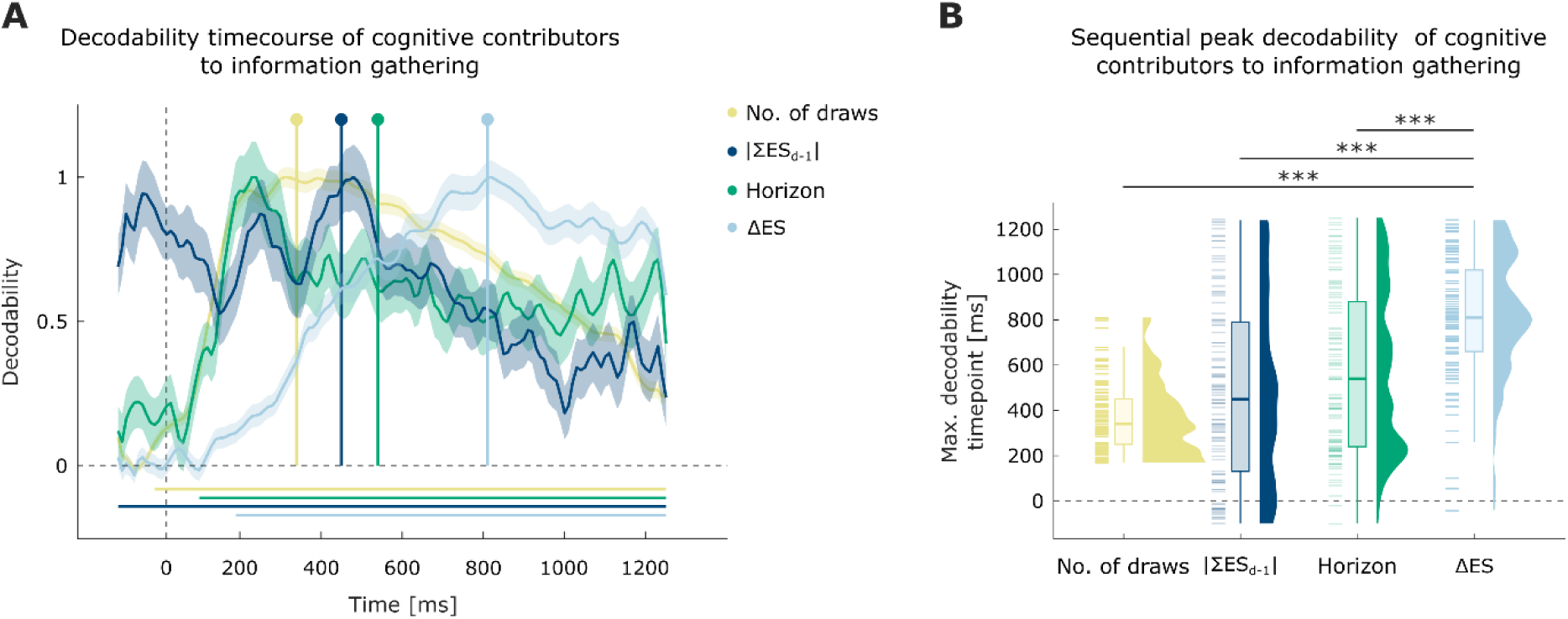
Sequential decodability of cognitive contributors to information gathering from whole-brain MEG activity. **A** Cognitive factors are predicted from MEG activity using lasso regression following an iterative method described previously^54^. The decoded variables of current number of draws, horizon condition, cumulative evidence strength from the start of the game until the previous draw |ΣESd-1|, and ΔES were all significantly decodable (p < 0.05, cluster-based permutation test at threshold t > 3; denoted by horizontal lines). Peak decodability time points, quantified as the group average of individual maximum values, are indicated by lollipops. Correlation values were normalised by dividing these by the maximum correlation value for visualization purposes. Note that all statistical tests were performed prior to normalisation. **B** Maximum decodability is achieved earlier in the trial for factors independent of the currently presented stimulus (current number of draws, horizon condition, the cumulative evidence from the start of the game until the previous draw) than for those dependent on current evidence processing (ΔES). Asterisks denote the level of significance (* p < 0.05, ** p < 0.01, *** p < 0.001).

*Reduced neural signatures of ΔES in high-OC participants mirror behavioural findings* Having established the decodability of cognitive contributors to information gathering, we next tested whether these representations were altered in high-OC participants. More concretely, given a reduced effect of ΔES on behaviour in the latter group, we hypothesised a reduced strength of this representation in the brain. On this basis, we assessed how transdiagnostic OC symptoms related to the decodability of ΔES. We found that ΔES representations were attenuated in high-OC participants, whereby two time periods reached significance using cluster-based permutation testing, namely 420-470 ms and 530-560 ms after stimulus onset (Fig. 4A-B; controlling for other psychiatric factors). Neither the decodability of the variable nor the individual differences appear driven by the effect of a commitment to a final decision, as both decodability and individual differences associated with the OC factor remained consistent after omitting the sample immediately preceding the response (see Supplementary Fig. 5A). Likewise, the results held for the cumulative evidence strength for the current majority ΣESd, as opposed to the update in evidence strength ΔES (see Supplementary Fig. 5B). When analysing other variables, we did not find any links with the OC spectrum. This means that – as per behavioural findings – the ΔES representation is attenuated in participants with high OC symptoms.

**Figure 4:**
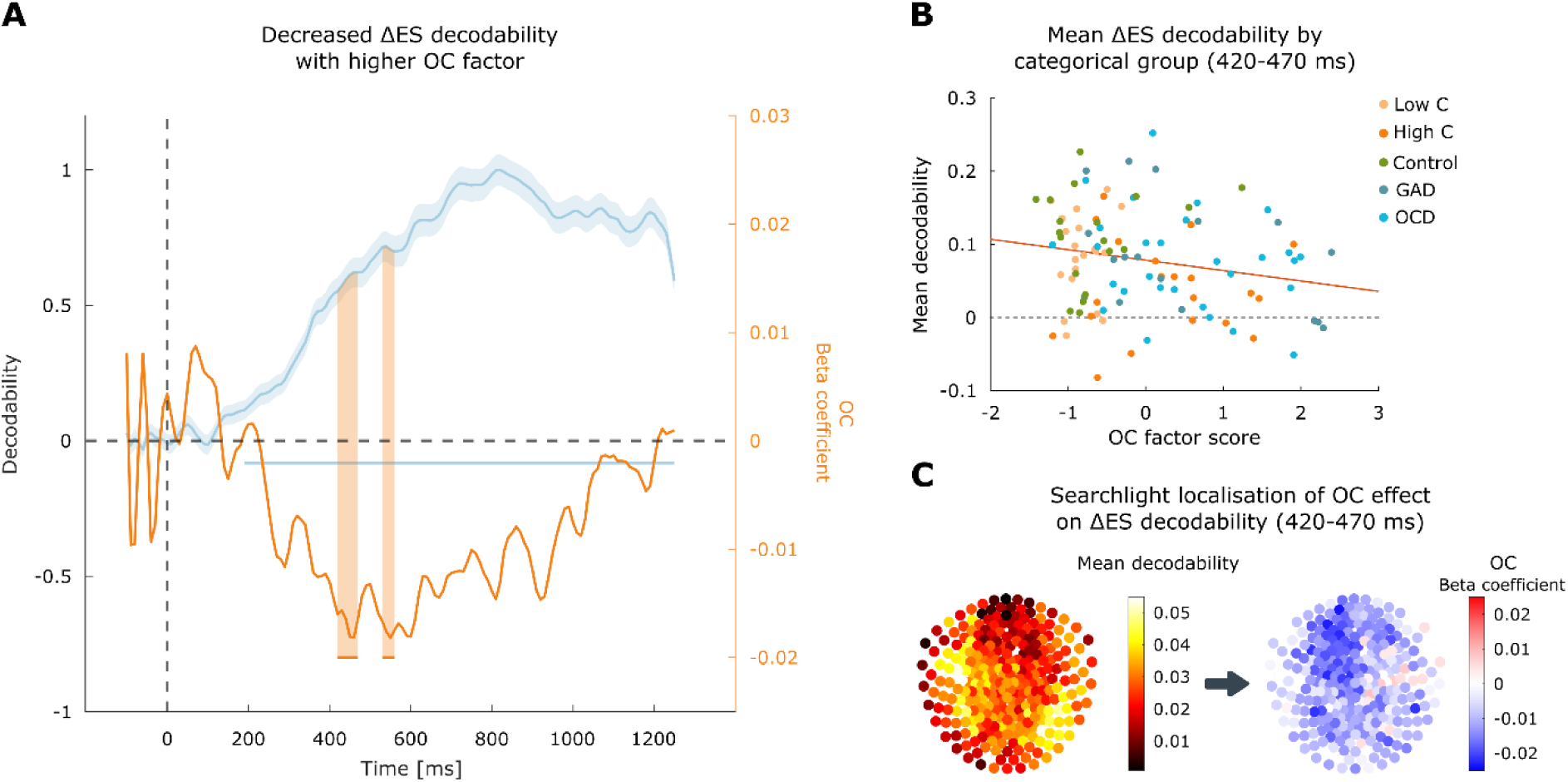
Reduced decodability of ΔES from MEG activity linked to OC symptoms. **A** ΔES can be decoded from MEG activity using an iterative multivariate lasso regression (left y-axis). Individual differences in OC factor predict significantly decreased decodability within 420-470 ms and 530-560 ms post stimulus (right y-axis). **B** Illustration of these attenuated ΔES effects within 420-470 ms. **C** To investigate the spatial distribution of the sensors contributing to the decodability of ΔES within the time windows where we find individual differences (420-470 ms post stimulus onset), we used searchlight analyses. ΔES are represented in a wider network comprising frontal, mediocentral and occipital areas (left). The attenuation of ΔES representations in high-OC participants was primarily driven by mediofrontal sensors, supporting the idea of altered ΔES processing in mediofrontal areas in high-OC participants (right).

### Attenuation of ΔES representations arise from mediofrontal areas

Lastly, we investigated the likely regional origins of an attenuated OC spectrum-related ΔES signal. While we found a relatively widespread activation pattern for ΔES per se (Fig. 4C, left panel), the attenuation in high-OC participants was pronounced in mediofrontal areas (Fig. 4C, right panel; see Supplementary Fig. 6 for the results for the second time window and Supplementary Figs. 7 and 8 for alternative methods). This suggests that the observed indecisiveness may arise from altered ΔES processing in mediofrontal areas.

## Discussion

Indecisiveness and doubt are common experiences in everyday life, but can reach debilitating extremes in mental health conditions, notably OCD. Some aspects of these symptoms can be quantitatively captured as excessive information gathering, yet their roots remain unclear^4,10,16,18,19,21–23^. Here, using the largest ever behavioural dataset in combination with data from a novel neuroimaging task, we show that attenuated weighting of most recent information by means of evidence-strength updates (ΔES) drives altered information gathering across an OC spectrum. Our finding of attenuated ΔES was evident at the level of both behaviour and brain, and was also evident dimensionally and transdiagnostically across both patients and those with non-clinical symptoms.

Excessive information gathering has been extensively investigated in relation to OCD and OC-like symptoms^4,10,16,18,19,21–23^, yet the precise mechanisms remain unclear. Much previous work on information gathering (including our own) has focused on altered decision criteria, such as collapsing decision thresholds or urgency signals^4,34^, ignoring the possibility that evidence integration (i.e., how information gathered at different timepoints is combined) might be implicated. We show that information integration shows a strong recency effect, where most recent information (by means of ΔES) exerts a much bigger impact on deciding, and when to decide, than more remote information. A recency weighting has been argued to confer an advantage under certain signal strength conditions^55^ or in dynamic contexts where the latest information is most representative^56^. It is also well aligned with similar recency biases found across domains that span perception, memory and mood ^37,38,57,58^. Interestingly, the nonlinear accrual of evidence has been widely accepted in the reward learning domain^59,60^, and the present results indicate similar mechanisms are at play also during information gathering. Crucially, ΔES is also represented in the brain, as a late electrophysiological component peaking approximately 920 ms after new information is revealed. Importantly, this ΔES representation peaks following the representation of other decision variables from which it is computed (specifically, newly revealed information and prior evidence), further supporting a putative sequential cascade of information processing^61,62^.

A mediofrontal localisation of the ΔES decodability is strongly reminiscent of that found in previous studies on information sampling during value-based decisions, where consistency of current evidence with a participant’s belief is signalled by anterior cingulate cortex (ACC) activation in single-cell recordings from monkeys^63^ and in an equivalent human fMRI paradigm^64^. In line with the strong effect of ΔES on the probability of committing to a decision, ACC signals peak immediately prior to the final decision in monkeys^63^. While the signal we show here does not depend crucially on the brain activity immediately prior to commitment to a decision (see Supplementary Fig. 5A), it should be noted that this possibility would be best addressed using a response-locked analysis, as opposed to the stimulus-locked approach we chose, and remains a question for future studies.

The ΔES measure may also fit within a broader belief-updating or predictive coding framework, provided the evidence strength for the current majority maps onto a participant’s current belief. If this assumption holds, ΔES can be cast as a type of prediction error (PE), in that it captures the deviation from the expectation formed in the previous draw. While different prediction errors have been described beyond the classic reward prediction errors, from perceptual to cognitive^57,65^, ΔES is different in so far as it is not based on explicit feedback for a completed action but rather on novel information available during the decision making process. As such, its relationship to a traditional conceptualisation of PEs needs to be established. This also applies to its implementation in the brain as well - while classic reward PEs have been associated with dopamine signalling, the extent to which dopamine influences information gathering is less clear^34,66,67^ and other candidate neurotransmitter systems, such as noradrenaline or serotonin, may conceivably contribute to the formation of ΔES^34,36^.

Critically for this study, we find attenuation of ΔES in people with (clinically and non-clinically) high OC symptoms and we propose it is the likely driver underlying excess information gathering behaviour found by draws-to-decision^4,10,18–23^ (though see also^16,17^). Importantly, we find a closer association between the ΔES weighting and the OC dimension, than with draws to a decision alone, supporting the notion this is the driving factor, while the latter behavioural marker is likely also under the influence of many other cognitive contributors, such as stringent constraints, urgency signals, and clear speed-accuracy trade-offs (as found in our MEG experiment). The finding of attenuated ΔES in OC, without a significant effect on draws to a decision, in the more constrained MEG experiment (cf. horizon conditions) also speaks to this, and accords with previous literature showing intact adjustments in speed-accuracy trade-offs in OCD patients^4,22^ Our finding of an attenuated neural representation of ΔES in high-OC participants also suggests that it may provide a sensitive biomarker. Indeed, the mediofrontal regions which appear most implicated in the attenuation of the ΔES representation yield a tantalising target for future investigations.

A diametrically opposite information gathering bias prominent in mental health disorders is jumping to conclusions (JTC), particularly seen in patients with psychosis^9,68–70^. Whether this JTC bias is driven by similar or distinct neurocognitive mechanisms remains unexplored. It is interesting, however, that similar imbalanced weighting hypotheses have been proposed for JTC^71–73^, along with the notion of altered PE processing in both psychosis and schizophrenia^74^.

Establishing a more refined marker of information gathering biases^40^ is likely to be crucial both for understanding indecisiveness and building new interventions. Indecisiveness manifests in different ways in the real world^23^ with different degrees of severity, which motivated the dimensional and transdiagnostic approach adopted here that included patients diagnosed with OCD and with generalised anxiety (GAD). We found that OC symptoms are not exclusive to OCD patients but are prominent in some GAD patients and non-clinical, high OC, participants in our sample (see Fig. 2B). While data on indecisiveness in clinically diagnosed GAD is lacking, our findings align with previous reports of high indecisiveness in undiagnosed individuals with probable GAD^75^ as well as in those with high levels of symptoms of anxiety and depression^76,77^. Moreover, our findings supports a transdiagnostic approach adopted in previous studies where dimensional measures outperformed traditional diagnostic categories^53^. Thus, we propose that indecisiveness and excessive information gathering is a transdiagnostic symptom and one that is not limited to OCD – indeed, indecisiveness in depression and dependent personality disorder has previously been highlighted^76^.

In sum, we provide evidence of an attenuated weighting of ΔES, both at the behavioural and neural level, linked to transdiagnostic OC symptoms, in a large crowd-sourced sample as well as a lab-based sample including clinical patients. This novel neurocognitive marker may bring us closer to understanding the mechanisms which drive excessive information gathering and indecisiveness across health and mental illnesses, and open avenues to the development of novel and improved interventions for these pervasive symptoms.

## Methods

We collected two types of data: a non-clinical large-scale sample completed a gamified information gathering task using the Brain Explorer app for hand-held devices as well as the Obsessive-Compulsive Inventory-Revised (OCI-R). A second in-person sample including OCD and GAD patients together with matched controls and low and high compulsive non-patients completed an analogous information gathering task while their MEG activity was recorded. In both studies, participants completed variants of information gathering tasks^78^ in which they were asked to indicate which of two possible stimuli was more plentiful. They had the choice to either continue sampling additional information or stop and make a binary choice between the two stimuli. We experimentally varied the amount of current and past information and the maximum sampling time before making a decision to assess their impact on information gathering and the relationship with OC symptoms.

### Participants

**Smartphone population sample**: At the time of analysis 8,670 participants completed the gamified information gathering task on Brain Explorer (www.brainexplorer.net), of whom 6,743 had also completed the OCI-R (data collected between November 2020 and August 2023). We analysed each participant’s first complete game (N = 15 trials) that fulfilled the inclusion criteria, which comprised a mean number of draws between 2 and 23, accuracy above 80%, and a minimum of three unique values in their number of draws to a decision, resulting in a final sample size of 5,237.

Additional exclusion criteria were applied to conduct the confidence analysis, namely a mean confidence rating between 15 and 98 and a minimum of three unique values in their confidence ratings, yielding a sample size of 4,302. Here, participants with any outlier beta coefficients in the GLM (see below), defined as values 1.5 times the interquartile range above the third quartile or below the first quartile, were removed, resulting in a final sample size of 3,293.

**In-lab MEG sample**: 115 participants completed the MEG study (data collected between April 2015 and August 2018). OCD and GAD patients were recruited through NHS services, charities and advertisements. Controls were recruited in areas of similar socioeconomic status as the patients’ to ensure comparable backgrounds. All participants underwent a structured clinical interview (SCID)^79^ conducted by experienced researchers. All participants with OCD and GAD fulfilled the corresponding diagnostic criteria for only one of these disorders^12^. In addition, self-reported symptom severity in patients was assessed using the Y-BOCS interview^15^. Exclusion criteria were current use of antipsychotic medication, severe learning disability, and comorbidities including current psychosis or bipolar disorder, autism spectrum disorder, substance abuse disorder, tics or Tourette disorder, and personality disorders except for obsessive-compulsive personality disorder.

Data from the high and low OC groups have been reported previously^80^. Briefly, participants were recruited from a large population-based sample of young people (U- CHANGE study; www.nspn.org.uk)^81,82^ based on the revised Padua Inventory – Washington State University Revision (PI-WSUR^83^).

Two participants were excluded because they did not meet the clinical criteria for GAD, one participant was excluded because of comorbid OCD and GAD, one was excluded due to fully missing questionnaire data. One additional participant was excluded due to an excessive number of non-decision trials (∼55%, compared to a mean of ∼20% across the remaining participants), resulting in below-chance performance. Five further participants were excluded due to technical difficulties and/or poor MEG data quality. A total of 105 participants were thus included in the analysis. The final groups comprised OCD patients (n = 29), GAD patients (n = 17), matched controls (n = 19), and non-clinical high compulsive (n = 20) and low compulsive groups (n = 20). Group demographics, medication percentages, and comorbidities for the final sample included in the analysis are reported in Supplementary Table 1.

The study was approved by the University College London (UCL) and NHS research ethics committees, and all participants gave written informed consent.

### Task

Participants in both studies completed an information gathering task where their goal was to determine which of two possible stimuli was more plentiful. Each draw consisted of a new sample of the stimuli, after which the participant made the decision to either continue sampling additional information or stop and make a binary choice between the two stimuli.

**Smartphone population sample**: During each game (Fig. 1A), participants viewed a grid of 25 locations, each of which could contain one of the two possible stimuli, here gems of different shape and colour. They could view which of two possible gems was hidden at a given position by tapping on it on the grid or make a binary choice indicating which of the two gems was more plentiful by tapping on the corresponding gem at the bottom of the screen. Uncovered gems remained on the screen until the participant made a binary choice. There were no constraints or penalties on the response time or the number of draws. After each decision, participants gave a confidence rating using a slider, and then received feedback on their performance in the form of a reward or a loss. They won 100 points for every correct choice and lost 100 points for every incorrect choice. Each session consisted of interactive instructions, which included a simplified mock game, followed by the 15 games which were included in the analysis.

**In-lab MEG sample**: To better disentangle cognitive contributing factors and their neural representations, we designed an entirely novel task optimised for MEG. In this version, the two possible products were always blue zircons or gold nuggets, represented by blue and yellow squares on a grey background respectively. Evidence samples consisted of 5 squares at a time which varied in their proportion of yellow to blue, and were presented automatically every 1,250 ms, i.e., participants did not actively gather information as in the Brain Explorer task. Participants were instructed that samples were recovered by a robot which may run out of batteries, such that here there was a variable finite number of draws per game, termed the horizon. The maximum number of draws varied between 4 and 8 in the short horizon condition and between 10 and 14 in the long horizon condition. These conditions were cued by a different coloured frame (green or pink, assigned randomly for each participant). There were 80 games in each horizon condition, randomly intermixed. Once participants committed to a choice, they made their response by pressing one of two buttons on a MEG-compatible button-box. The length of the draw sequence was presented in full independently of when the participant made their decision – in other words, if the participant made a decision before the last draw, the draw sequence continued until it reached its pre-determined end. Participants received thorough instructions and completed 12 games as part of a practice block which included the horizon manipulation prior to beginning the experiment, such that they acquired a good intuition about the horizon length already during the practice block. They received 2 points for a correct response, lost 2 points for an incorrect response, and lost 1 point for non-decisions. They could track their performance and corresponding rewards by means of a blue bar at the bottom of the screen, which started at the 50% mark. They were told that each time the bar reached the maximum threshold of 100%, they would receive an additional £0.50 at the end of the experiment.

### OC symptoms assessment

**Smartphone population sample**: Participants completed the OCI-R^48^ within the Brain Explorer app.

**In-lab MEG sample**: Participants completed seven self-report questionnaires assessing psychiatric symptoms: Barratt Impulsiveness Scale (BIS^84^), Beck Depression Inventory II (BDI-II^85^), Intolerance of Uncertainty Scale (IUS^86^), OCI-R^48^, PI-WSUR^83^, State and Trait Anxiety Inventory (STAI^87^), and Frost Multidimensional Perfectionism Scale (FMPS^88^). Participants additionally completed the matrix and vocabulary subtests of the Wechsler Abbreviated Scale of Intelligence (WASI^89^) and the Edinburgh inventory^90^ was collected to assess handedness. Missing responses in the questionnaires were imputed with the mean response, provided the number of missing items did not exceed 20% of the items in the corresponding scale.

We conducted an exploratory factor analysis (EFA) on the 210 individual items which compose the seven psychiatric symptom questionnaires to reduce the dimensionality of the data. The EFA solution was estimated using maximum likelihood and oblimin rotation based on the heterogeneous correlation matrix to account for both continuous and binary correlations using the *psych* package in R^91^. The number of factors was determined based on the Cattell-Nelson-Gorsuch index^92^, i.e., the eigenvalue at which the greatest difference between two successive slopes occurs, visible as a sharp elbow in a scree plot, implemented with the *nFactors* package in R^93^. Although the sample size resulted in a relatively low ratio of observations to items, adequate sample size for factor analyses ultimately depends on the correlation structure in the data. Indeed, in this case the Kaiser-Meyer-Olkin (KMO) Measure of Sampling Adequacy was 0.94 and Bartlett’s Test of Sphericity was significant (p < 0.001), in line with published recommendations^94^. More detailed results of the EFA can be found in the Supplementary Materials.

### Behavioural analysis

**Smartphone population sample**: We correlated the primary index of information gathering, namely the number of draws to a decision, as well as the accuracy with the OCI-R scores using Spearman’s correlations.

We then investigated the underlying processes of information gathering by fitting participants’ draw-wise binary choice to sample information or not to a logit-linked binomial general linear mixed model (GLMM) predicting the probability to make a decision from the experimental factors, OCI-R scores, and the interactions of OCI-R scores with the experimental factors, as well as random intercepts and slopes for each experimental factor. Experimental factors consisted of, first, the cumulative evidence strength on the previous draw

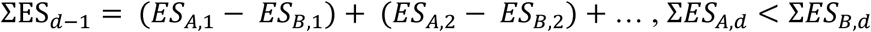

i.e., the total number of samples in favour of stimulus A across all draws ΣESA until draw d-1 minus the total number of samples in favour of stimulus B across all draws ΣESB until draw d-1 for the current majority, second, the change in evidence strength on the current draw or evidence strength updater (ΔES)

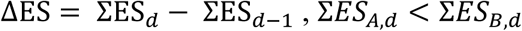

i.e., the difference in relative evidence in draw d ΣESd and the previous draw |ΣESd-1|, and, third, the current draw number.

Lastly, we conducted a mediation analysis evaluating the relationship between confidence, OCI-R and sensitivity to current evidence using the M3 mediation toolbox in MATLAB^51,52^. To perform this analysis, we obtained a measure of sensitivity to ΔES for each individual participant by fitting a separate general linear model (GLM) to each participant predicting the probability to make a decision, based on the same experimental factors as in the GLMM. We then tested if the individual beta coefficients for ΔES mediated the negative association between mean confidence and OCI-R scores.

**In-lab MEG sample**: We performed an analogous analysis, while accounting for sample and task differences. As stated above, we used the OC factor score to quantify variability in OC symptoms. In addition, we adapted the GLMM by including the horizon condition and the interactions between the horizon and all other regressors. The interaction of ΔES with the horizon was omitted in the participant-level GLMs, as the beta coefficient was not significantly different from zero.

### MEG analysis

#### Data acquisition and preprocessing

Participants sat upright in the scanner while MEG was recorded continuously using a 275-channel system, whereby sensor MLO42 was missing for all participants, and in addition sensor MRC12 was missing for 59 participants. Empty channels were added in these cases in order to keep the dimensionality of the MEG data consistent across participants. Bad-quality channels were removed and not interpolated. Physiological and technical artefact removal was performed manually using Fieldtrip. The data was down-sampled to 100 Hz and high-pass filtered at 0.5 Hz to remove slow drift effects. Data was segmented into 1350-ms segments from -100 to 1250ms relative to stimulus onset. MEG data was z-scored by block and channel (univariate normalisation).

#### Decoding analysis

Whole-brain MEG activity was used to decode ΔES using an iterative multivariate lasso regression^54^. Here, we estimated the lasso regression 2,000 times, using a random subset of 50 sensors on each iteration. The primary benefit of this method is the increased reliability of the individual differences analysis, relative to a single iteration of a multivariate lasso regression using all channels. The regularization parameter λ was set at 0.01 using 5-fold cross-validation. The value for the regularization parameter was tuned by optimizing the cross-validation performance of an orthogonal variable, namely the current evidence for yellow (correlation between current evidence for yellow and ΔES: r = 2.755 × 10^-4^, p = 0.920), and was fixed for all subsequent whole-brain analyses. The initial optimization procedure was conducted using a non-iterative multivariate lasso regression using all 273 channels, yet note that the decodability using the iterative method remains extremely similar for a range of regularization parameter values in follow up analyses (see Supplementary Fig. 4D).

The correlation between the tested and trained data constituted the measure of decodability. Non-parametric cluster-based permutation tests against 0 were used to determine the time window at which the decodability was significant, setting the t statistic threshold at 3 and using one-sided p < 0.05. Individual differences in decodability were assessed by predicting the decodability at each time point from the three factor scores in a GLM per time point. Significant differences were assessed by non-parametric cluster-based permutation tests at each time point in which the factor scores were shuffled across participants while maintaining the factor score combination constant (t = 2, p < 0.025, two-sided test). Correlation values were normalized by dividing these by the maximum correlation value for visualization purposes, yet note that all statistical tests were performed prior to normalisation.

To investigate which brain areas underpin the decoding, we used a searchlight analysis, such that the lasso regression was estimated 273 times, i.e., for each sensor and its direct neighbors, determined using the Fieldtrip *ft_prepare_neighbours* function. Given the stark difference in the number of sensors used in the searchlight analysis, we first fine-tuned the value of the regularization parameter λ in the same way as above, and set it at 0.001. We then calculated the mean decodability per sensor within a given time window of interest to generate sensor contribution maps. To explore individual differences in the sensor contribution maps, we fit a GLM predicting the sensor contribution per sensor averaged across the time window of interest from the three factor scores. In order to keep the sensor subsets constant across participants, sensors MLO42 and MRC12 were omitted for all.

## Acknowledgements

We thank the Central London OCD support group, the OCD Action charity, the North East London NHS foundation, especially the IAPT services at Waltham Forest, Redbridge and Barking & Dagenham, and the Camden and Islington NHS talking therapies service iCope for their help with patient recruitment. TUH is supported by a Sir Henry Dale Fellowship (211155/Z/18/Z; 211155/Z/18/B; 224051/Z/21) from Wellcome & Royal Society, a grant from the Jacobs Foundation (2017- 1261-04), the Medical Research Foundation, a 2018 NARSAD Young Investigator grant (27023) from the Brain & Behavior Research Foundation, and a Philip Leverhulme Prize from the Leverhulme Trust (PLP-2021-040). TUH is also supported by the Carl-Zeiss-Stiftung. A Wellcome Trust Cambridge–UCL Mental Health and Neurosciences Network grant (095844/Z/11/Z) supported this work. The Max Planck UCL Centre is a joint initiative supported by UCL and the Max Planck Society. The Wellcome Centre for Human Neuroimaging is supported by core funding from the Wellcome Trust (203147/Z/16/Z). TUH consults for Limbic Ltd. and holds shares in the company, which is unrelated to the current project. All other authors declare no conflicts of interest. This research was funded in whole, or in part, by the Wellcome Trust (211155/Z/18/Z). For the purpose of Open Access, the author has applied a CC BY public copyright license to any Author Accepted Manuscript version arising from this submission.

## References

1. Bogacz, R., Wagenmakers, E.-J., Forstmann, B. U. & Nieuwenhuis, S. The neural basis of the speed–accuracy tradeoff. Trends Neurosci. 33, 10–16 (2010).

2. Schouten, J. F. & Bekker, J. A. M. Reaction time and accuracy. Acta Psychol. (Amst.) 27, 143–153 (1967).

3. Bogacz, R., Hu, P. T., Holmes, P. J. & Cohen, J. D. Do humans produce the speed–accuracy trade-off that maximizes reward rate? Q. J. Exp. Psychol. 63, 863–891 (2010).

4. Hauser, T. U. et al. Increased decision thresholds enhance information gathering performance in juvenile Obsessive-Compulsive Disorder (OCD). PLOS Comput. Biol. 13, e1005440 (2017).

5. Phillips, L. D. & Edwards, W. Conservatism in a simple probability inference task. (1966).

6. Hemsley, D. R. & Garety, P. A. The Formation of Maintenance of Delusions: a Bayesian Analysis. Br. J. Psychiatry 149, 51–56 (1986).

7. Moritz, S. et al. A two-stage cognitive theory of the positive symptoms of psychosis. Highlighting the role of lowered decision thresholds. J. Behav. Ther. Exp. Psychiatry 56, 12–20 (2017).

8. Ward, T. & Garety, P. A. Fast and slow thinking in distressing delusions: A review of the literature and implications for targeted therapy. Schizophr. Res. 203, 80–87 (2019).

9. McLean, B. F., Mattiske, J. K. & Balzan, R. P. Association of the Jumping to Conclusions and Evidence Integration Biases With Delusions in Psychosis: A Detailed Meta-analysis. Schizophr. Bull. 43, 344–354 (2017).

10. Fear, C. F. & Healy, D. Probabilistic reasoning in obsessive–compulsive and delusional disorders. Psychol. Med. 27, 199–208 (1997).

11. Koutsouleris, N., Hauser, T. U., Skvortsova, V. & Choudhury, M. D. From promise to practice: towards the realisation of AI-informed mental health care. *Lancet Digit*. Health 4, e829–e840 (2022).

12. Diagnostic and Statistical Manual of Mental Disorders: DSM-5. (American Psychiatric Association, Washington, D.C, 2013).

13. The ICD-10 Classification of Mental and Behavioural Disorders: Clinical Descriptions and Diagnostic Guidelines. (World Health Organization, Geneva, 1992).

14. Taillefer, S. E., Liu, J. J. W., Ornstein, T. J. & Vickers, K. Indecisiveness as a predictor of quality of life in individuals with obsessive and compulsive traits. J. Obsessive-Compuls. Relat. Disord. 10, 91–98 (2016).

15. Storch, E. A. et al. Development and psychometric evaluation of the Yale–Brown Obsessive-Compulsive Scale—Second Edition. Psychol. Assess. 22, 223–232 (2010).

16. Grassi, G. et al. Think twice: Impulsivity and decision making in obsessive–compulsive disorder. (2015) doi:10.1556/2006.4.2015.039.

17. Jacobsen, P., Freeman, D. & Salkovskis, P. Reasoning bias and belief conviction in obsessive-compulsive disorder and delusions: Jumping to conclusions across disorders? Br. J. Clin. Psychol. 51, 84–99 (2012).

18. Pélissier, M.-C. & O’Connor, K. P. Deductive and inductive reasoning in obsessive-compulsive disorder. Br. J. Clin. Psychol. 41, 15–27 (2002).

19. Volans, P. J. Styles of Decision-making and Probability Appraisal in Selected Obsessional and Phobic Patients. Br. J. Soc. Clin. Psychol. 15, 305–317 (1976).

20. Voon, V. et al. Decisional impulsivity and the associative-limbic subthalamic nucleus in obsessive-compulsive disorder: stimulation and connectivity. Brain 140, 442–456 (2017).

21. Erhan, C. et al. Disrupted latent decision processes in medication-free pediatric OCD patients. J. Affect. Disord. 207, 32–37 (2017).

22. Banca, P. et al. Evidence Accumulation in Obsessive-Compulsive Disorder: the Role of Uncertainty and Monetary Reward on Perceptual Decision-Making Thresholds. Neuropsychopharmacology 40, 1192–1202 (2015).

23. Loosen, A. M., Skvortsova, V. & Hauser, T. U. Obsessive–compulsive symptoms and information seeking during the Covid-19 pandemic. Transl. Psychiatry 11, 1–10 (2021).

24. Cisek, P. & Thura, D. Models of Decision-Making Over Time. in *Oxford Research Encyclopedia of Neuroscience* (Oxford University Press, 2022). doi:10.1093/acrefore/9780190264086.013.346.

25. Gold, J. I. & Shadlen, M. N. The Neural Basis of Decision Making. Annu. Rev. Neurosci. 30, 535–574 (2007).

26. Brody, C. D. & Hanks, T. D. Neural underpinnings of the evidence accumulator. Curr. Opin. Neurobiol. 37, 149–157 (2016).

27. Shadlen, M. N. & Shohamy, D. Decision making and sequential sampling from memory. Neuron 90, 927–939 (2016).

28. Krajbich, I., Armel, C. & Rangel, A. Visual fixations and the computation and comparison of value in simple choice. Nat. Neurosci. 13, 1292–1298 (2010).

29. Busemeyer, J. R. & Townsend, J. T. Decision field theory: A dynamic-cognitive approach to decision making in an uncertain environment. Psychol. Rev. 100, 432–459 (1993).

30. Cisek, P., Puskas, G. A. & El-Murr, S. Decisions in changing conditions: The urgency-gating model. J. Neurosci. 29, 11560–11571 (2009).

31. Drugowitsch, J., Moreno-Bote, R., Churchland, A. K., Shadlen, M. N. & Pouget, A. The Cost of Accumulating Evidence in Perceptual Decision Making. J. Neurosci. 32, 3612– 3628 (2012).

32. Milosavljevic, M., Malmaud, J., Huth, A., Koch, C. & Rangel, A. The Drift Diffusion Model can account for the accuracy and reaction time of value-based choices under high and low time pressure. Judgm. Decis. Mak. 5, 437–449 (2010).

33. Thura, D., Beauregard-Racine, J., Fradet, C.-W. & Cisek, P. Decision making by urgency gating: theory and experimental support. J. Neurophysiol. 108, 2912–2930 (2012).

34. Hauser, T. U., Moutoussis, M., Purg, N., Dayan, P. & Dolan, R. J. Beta-Blocker Propranolol Modulates Decision Urgency During Sequential Information Gathering. J. Neurosci. 38, 7170–7178 (2018).

35. Bowler, A. et al. Children perform extensive information gathering when it is not costly. Cognition 208, 104535 (2021).

36. Michely, J., Martin, I. M., Dolan, R. J. & Hauser, T. U. Boosting Serotonin Increases Information Gathering by Reducing Subjective Cognitive Costs. J. Neurosci. 43, 5848– 5855 (2023).

37. Wyart, V., Myers, N. E. & Summerfield, C. Neural Mechanisms of Human Perceptual Choice Under Focused and Divided Attention. J. Neurosci. 35, 3485–3498 (2015).

38. Cheadle, S. et al. Adaptive Gain Control during Human Perceptual Choice. Neuron 81, 1429–1441 (2014).

39. Osth, A. F. & Farrell, S. Using response time distributions and race models to characterize primacy and recency effects in free recall initiation. Psychol. Rev. 126, 578–609 (2019).

40. Carland, M. A., Thura, D. & Cisek, P. The urge to decide and act: implications for brain function and dysfunction. The Neuroscientist 25, 491–511 (2019).

41. Bogacz, R. Optimal decision-making theories: linking neurobiology with behaviour. Trends Cogn. Sci. 11, 118–125 (2007).

42. Hogarth, R. M. & Einhorn, H. J. Order effects in belief updating: The belief-adjustment model. Cognit. Psychol. 24, 1–55 (1992).

43. Usher, M. & McClelland, J. L. The time course of perceptual choice: The leaky, competing accumulator model. Psychol. Rev. 108, 550–592 (2001).

44. Bogacz, R., Brown, E., Moehlis, J., Holmes, P. & Cohen, J. D. The physics of optimal decision making: A formal analysis of models of performance in two-alternative forced-choice tasks. Psychol. Rev. 113, 700–765 (2006).

45. Hauser, T. U. et al. Increased fronto-striatal reward prediction errors moderate decision making in obsessive–compulsive disorder. Psychol. Med. 47, 1246–1258 (2017).

46. Moutoussis, M., Bentall, R. P., El-Deredy, W. & Dayan, P. Bayesian modelling of Jumping-to-Conclusions bias in delusional patients. Cognit. Neuropsychiatry 16, 422–447 (2011).

47. Huq, S. F., Garety, P. A. & Hemsley, D. R. Probabilistic Judgements in Deluded and Non- Deluded Subjects. Q. J. Exp. Psychol. Sect. A 40, 801–812 (1988).

48. Foa, E. B. et al. The Obsessive-Compulsive Inventory: Development and validation of a short version. Psychol. Assess. 14, 485–496 (2002).

49. Hauser, T. U., Moutoussis, M., Dayan, P. & Dolan, R. J. Increased decision thresholds trigger extended information gathering across the compulsivity spectrum. Transl. Psychiatry 7, 1–10 (2017).

50. Dar, R., Sarna, N., Yardeni, G. & Lazarov, A. Are people with obsessive-compulsive disorder under-confident in their memory and perception? A review and meta-analysis. Psychol. Med. 52, 2404–2412.

51. Wager, T. D., Davidson, M. L., Hughes, B. L., Lindquist, M. A. & Ochsner, K. N. Prefrontal-subcortical pathways mediating successful emotion regulation. Neuron 59, 1037–1050 (2008).

52. Wager, T. D. et al. Brain mediators of cardiovascular responses to social threat: Part I: Reciprocal dorsal and ventral sub-regions of the medial prefrontal cortex and heart-rate reactivity. NeuroImage 47, 821–835 (2009).

53. Gillan, C. M. et al. Comparison of the Association Between Goal-Directed Planning and Self-reported Compulsivity vs Obsessive-Compulsive Disorder Diagnosis. JAMA Psychiatry 77, 77–85 (2020).

54. Kurth-Nelson, Z., Barnes, G., Sejdinovic, D., Dolan, R. & Dayan, P. Temporal structure in associative retrieval. eLife 4, e04919 (2015).

55. Summerfield, C. & Parpart, P. Normative Principles for Decision-Making in Natural Environments. Annu. Rev. Psychol. 73, 53–77 (2022).

56. van Bergen, R. S. & Jehee, J. F. M. Probabilistic Representation in Human Visual Cortex Reflects Uncertainty in Serial Decisions. http://biorxiv.org/lookup/doi/10.1101/671958 (2019) doi:10.1101/671958.

57. Kiyonaga, A., Scimeca, J. M., Bliss, D. P. & Whitney, D. Serial Dependence across Perception, Attention, and Memory. Trends Cogn. Sci. 21, 493–497 (2017).

58. Eldar, E., Rutledge, R. B., Dolan, R. J. & Niv, Y. Mood as Representation of Momentum. Trends Cogn. Sci. 20, 15–24 (2016).

59. Rescorla, R. A. & Wagner, A. R. A theory of Pavlovian conditioning : Variations in the effectiveness of reinforcement and non-reinforcement. Class. Cond. Curr. Res. Theory 2, 64–69 (1972).

60. Emanuel, A. & Eldar, E. Emotions as computations. Neurosci. Biobehav. Rev. 144, 104977 (2023).

61. Parés-Pujolràs, E., Kelly, S. P. & Murphy, P. R. Dissociable encoding of evolving beliefs and momentary belief updates in distinct neural decision signals. Preprint at 10.1101/2024.05.15.594345 (2024).

62. Diaconescu, A. O. et al. Hierarchical prediction errors in midbrain and septum during social learning. Soc. Cogn. Affect. Neurosci. 12, 618–634 (2017).

63. Hunt, L. T. et al. Triple dissociation of attention and decision computations across prefrontal cortex. Nat. Neurosci. 21, 1471–1481 (2018).

64. Kaanders, P., Nili, H., O’Reilly, J. X. & Hunt, L. Medial Frontal Cortex Activity Predicts Information Sampling in Economic Choice. J. Neurosci. 41, 8403–8413 (2021).

65. Den Ouden, H. E., Kok, P. & De Lange, F. P. How Prediction Errors Shape Perception, Attention, and Motivation. Front. Psychol. 3, (2012).

66. Ersche, K. D. et al. Peripheral biomarkers of cognitive response to dopamine receptor agonist treatment. Psychopharmacology (Berl.) 214, 779–789 (2011).

67. Ermakova, A. O., Ramachandra, P., Corlett, P. R., Fletcher, P. C. & Murray, G. K. Effects of Methamphetamine Administration on Information Gathering during Probabilistic Reasoning in Healthy Humans. PLOS ONE 9, e102683 (2014).

68. Dudley, R., Taylor, P., Wickham, S. & Hutton, P. Psychosis, Delusions and the “Jumping to Conclusions” Reasoning Bias: A Systematic Review and Meta-analysis. Schizophr. Bull. 42, 652–665 (2016).

69. Ross, R. M., McKay, R., Coltheart, M. & Langdon, R. Jumping to Conclusions About the Beads Task? A Meta-analysis of Delusional Ideation and Data-Gathering. Schizophr. Bull. 41, 1183–1191 (2015).

70. Fine, C., Gardner, M., Craigie, J. & Gold, I. Hopping, skipping or jumping to conclusions? Clarifying the role of the JTC bias in delusions. Cognit. Neuropsychiatry 12, 46–77 (2007).

71. Ashinoff, B. & Horga, G. Evidence-Order Effects in Probabilistic Inference: Recency Bias and Delusion-Like Ideation Across the Psychosis Continuum. Biol. Psychiatry 87, S392– S393 (2020).

72. Scheunemann, J., Fischer, R. & Moritz, S. Probing the Hypersalience Hypothesis—An Adapted Judge-Advisor System Tested in Individuals With Psychotic-Like Experiences. Front. Psychiatry 12, 612810 (2021).

73. Ashinoff, B. K., Buck, J., Woodford, M. & Horga, G. The effects of base rate neglect on sequential belief updating and real-world beliefs. PLOS Comput. Biol. 18, e1010796 (2022).

74. Sterzer, P. et al. The Predictive Coding Account of Psychosis. Biol. Psychiatry 84, 634– 643 (2018).

75. Koerner, N., Mejia, T. & Kusec, A. What’s in a name? Intolerance of uncertainty, other uncertainty-relevant constructs, and their differential relations to worry and generalized anxiety disorder*. Cogn. Behav. Ther. 46, 141–161 (2017).

76. Rassin, E., Muris, P., Franken, I., Smit, M. & Wong, M. Measuring General Indecisiveness. J. Psychopathol. Behav. Assess. 29, 60–67 (2007).

77. Lauderdale, S. A. & Oakes, K. Factor Structure of the Revised Indecisiveness Scale and Association with Risks for and Symptoms of Anxiety, Depression, and Attentional Control. J. Ration.-Emotive Cogn.-Behav. Ther. 39, 256–284 (2021).

78. Clark, L., Robbins, T. W., Ersche, K. D. & Sahakian, B. J. Reflection Impulsivity in Current and Former Substance Users. Biol. Psychiatry 60, 515–522 (2006).

79. First, M. B. Structured Clinical Interview for the (SCID). in *The Encyclopedia of Clinical Psychology* 1–6 (John Wiley & Sons, Ltd, 2015). doi:10.1002/9781118625392.wbecp351.

80. Hauser, T. U., Allen, M., Rees, G. & Dolan, R. J. Metacognitive impairments extend perceptual decision making weaknesses in compulsivity. Sci. Rep. 7, 6614 (2017).

81. Whitaker, K. J. et al. Adolescence is associated with genomically patterned consolidation of the hubs of the human brain connectome. Proc. Natl. Acad. Sci. 113, 9105–9110 (2016).

82. Vértes, P. E. et al. Gene transcription profiles associated with inter-modular hubs and connection distance in human functional magnetic resonance imaging networks. Philos. Trans. R. Soc. B Biol. Sci. 371, 20150362 (2016).

83. Burns, G. L., Keortge, S. G., Formea, G. M. & Sternberger, L. G. Revision of the Padua Inventory of obsessive compulsive disorder symptoms: Distinctions between worry, obsessions, and compulsions. Behav. Res. Ther. 34, 163–173 (1996).

84. Stanford, M. S. et al. Fifty years of the Barratt Impulsiveness Scale: An update and review. Personal. Individ. Differ. 47, 385–395 (2009).

85. Beck, A. T., Steer, R. A., Ball, R. & Ranieri, W. F. Comparison of Beck Depression Inventories-IA and-II in Psychiatric Outpatients. J. Pers. Assess. 67, 588–597 (1996).

86. Buhr, K. & Dugas, M. J. The intolerance of uncertainty scale: psychometric properties of the English version. Behav. Res. Ther. 40, 931–945 (2002).

87. 87. Spielberger, C., Gorsuch, R., Lushene, R., Vagg, P. & Jacobs, G. Manual for the State-Trait Anxiety Inventory (Form Y1 – Y2). Palo Alto, CA: Consulting Psychologists Press; vol. IV (1983).

88. Frost, R. O., Marten, P., Lahart, C. & Rosenblate, R. The dimensions of perfectionism. Cogn. Ther. Res. 14, 449–468 (1990).

89. Wechsler, D. Wechsler Abbreviated Scale of Intelligence--Second Edition. (2011) doi:10.1037/t15171-000.

90. Oldfield, R. C. The assessment and analysis of handedness: The Edinburgh inventory. Neuropsychologia 9, 97–113 (1971).

91. Revelle, W. psych: Procedures for Psychological, Psychometric, and Personality Research. (2024).

92. Cattell, R. B. The Scree Test For The Number Of Factors. Multivar. Behav. Res. 1, 245– 276 (1966).

93. Raîche, G., Walls, T. A., Magis, D., Riopel, M. & Blais, J.-G. Non-Graphical Solutions for Cattell’s Scree Test. Methodology 9, 23–29 (2013).

94. Williams, B., Onsman, A. & Brown, T. Exploratory factor analysis: A five-step guide for novices. Australas. J. Paramed. 8, (2010).

