## Supplementary Materials for "Indecision and recency-weighted evidence integration in non-clinical and clinical settings"

del Río M[1,2], Trudel N[1,2], Prabhu G[2], Hunt LT[3], Moutoussis M[2], Dolan RJ[1,2,4]* & Hauser TU[1,2,5,6]*

[1]Max Planck UCL Centre for Computational Psychiatry and Ageing Research, University College London, London, United Kingdom

[2]Wellcome Centre for Human Neuroimaging, University College London, London, United Kingdom

[3]Department of Experimental Psychology, University of Oxford, Oxford, United Kingdom

[4]State Key Laboratory of Cognitive Neuroscience and Learning, IDG/McGovern Institute for Brain Research, Beijing Normal University, Beijing, China

[5]Department of Psychiatry and Psychotherapy, Faculty of Medicine, University of Tübingen, Tübingen, Germany

[6]German Centre for Mental Health (DZPG), Tübingen, Germany

* Shared senior authors

*Recency effect in information gathering*

In order to test specifically for a recency effect in the decision to commit during information gathering, we fit an additional GLMM predicting *p(decide)* from the current evidence at draw d, d-1 and d-2 relative to the chosen option. This showed a decreasing contribution of current evidence for lagged draws, both in the smartphone population sample ($ES_d$: β = 0.648 SE = 0.007, p < 0.001; $ES_{d-1}$: β = 0.253 SE = 0.004, p < 0.001; $ES_{d-2}$: β = 0.195 SE = 0.005, p < 0.001, see Supplementary Fig. 1A) and in the in-lab MEG sample ($ES_d$: β = 0.519 SE = 0.015, p < 0.001; $ES_{d-1}$: β = 0.212 SE = 0.013, p < 0.001; $ES_{d-2}$: β = 0.216 SE = 0.010, p < 0.001, see Supplementary Fig. 1B).

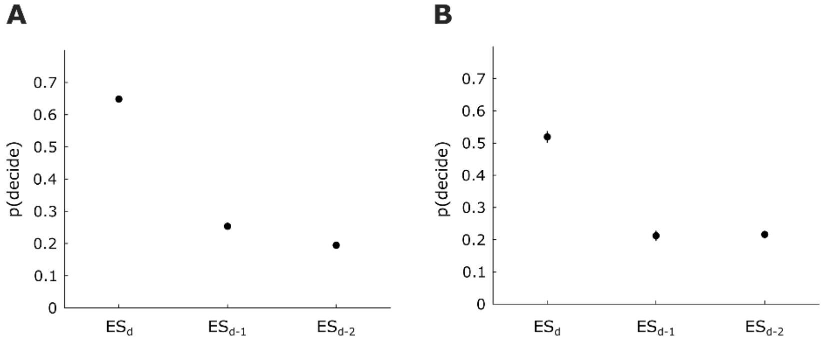


*Supplementary Figure 1: The probability of making a decision p(decide) was predicted from the current evidence at draw d, d-1, and d-2 relative to the chosen option. This showed a decreasing contribution of the evidence for lagged draws in **A** the smartphone population sample and **B** the in-lab MEG sample.*

*Self-reported OCD diagnosis effects on information gathering in smartphone population sample*

At enrolment in the smartphone population study, participants were prompted to self-report on their mental health history on a voluntary basis. We were thus able to compare those who self-reported a current diagnosis of OCD and those who self-reported no lifetime psychiatric disorder on the two main measures of interest, the number of draws to a decision and the weight of ΔES in a GLMM, where current OCD diagnosis was coded as a binary variable. We replicate the effect of increased number of draws to a decision in those who self-reported having a current diagnosis of OCD (N = 129) vs no lifetime psychiatric disorder (N = 3,948), albeit substantially less robustly (t(4075) = 1.655, p = 0.049, one-sided test). Likewise, the attenuation of the ΔES weight was not as robust and the attenuation of the weight of the previous evidence $|\Sigma ES_{d-1}|$ was not significant in those with self-reported OCD (OCD × $|\Sigma ES_{d-}$

$_1|$: $\beta = -0.058$, SE = 0.043, $p = 0.093$; OCD × ΔES: $\beta = -0.117$, SE = 0.060, $p = 0.026$, one-sided tests).

*In-lab MEG sample characterisation*

There were no significant differences in age, IQ or female/male ratio across the OCD, GAD and the control groups, assessed by means of paired t-tests or Fisher’s exact tests, as appropriate, for all combinations. The proportion of currently medicated patients did not quite reach significance when comparing OCD and GAD patients ($\chi^2(1, 46) = 3.669$, $p = 0.055$, as per a Pearson's Chi-squared test with Yates' continuity correction). The proportion of currently medicated OCD and GAD patients differed significantly from the control group as per Fisher’s exact tests ($p = 0.007$ and $p < 0.001$, respectively). The proportion of agoraphobia, body dysmorphic disorder (BDD), compulsive skin picking (CSP), trichotillomania, and social anxiety disorder (SAD) comorbidity did not differ significantly between OCD and GAD patients or between OCD/GAD patients and controls (all $p > 0.369$ as per Fisher’s exact tests). The proportion of depression was significantly higher relative to controls both for GAD ($p = 0.002$) and OCD patients ($p = 0.015$), but did not differ between OCD and GAD patients ($\chi^2(1, 46) = 0.388$, $p = 0.533$, as per a Pearson's Chi-squared test with Yates' continuity correction).

*Table 2: Participant characteristics: Age, sex ratio, IQ, comorbidities and medication per group, and Y-BOCS symptom severity scores for clinically diagnosed OCD patients.*

| | **Low Compulsive**, N = 20[1] | **High Compulsive**, N = 20[1] | **Control**, N = 19[1] | **OCD**, N = 29[1] | **GAD**, N = 17[1] |
|---|---|---|---|---|---|
| Age | 21.4 ± 2.5, (range 18.0 - 26.0) | 20.8 ± 2.3, (range 18.0 - 26.0) | 29.6 ± 8.4, (range 18.0 - 44.0) | 29.1 ± 8.5, (range 18.0 - 45.0) | 31.8 ± 8.7, (range 20.0 - 48.0) |
| Sex | | | | | |
| Male | 7 / 20 | 6 / 20 | 5 / 19 | 7 / 29 | 5 / 17 |
| Female | 13 / 20 | 14 / 20 | 14 / 19 | 22 / 29 | 12 / 17 |
| IQ | 113.7 ± 9.7 | 113.5 ± 8.7 | 111.4 ± 10.3 | 106.4 ± 14.3 | 112.0 ± 8.9 |
| Symptom severity YBOCS (total) | | | | 20.9 ± 6.0 | |
| Agoraphobia (%) | 0 / 20 (0%) | 0 / 20 (0%) | 0 / 19 (0%) | 0 / 29 (0%) | 1 / 17 (5.9%) |
| BDD (%) | 0 / 20 (0%) | 0 / 20 (0%) | 0 / 19 (0%) | 2 / 29 (6.9%) | 0 / 17 (0%) |
| CSP (%) | 0 / 20 (0%) | 0 / 20 (0%) | 0 / 19 (0%) | 1 / 29 (3.4%) | 0 / 17 (0%) |
| Depression (%) | 0 / 20 (0%) | 0 / 20 (0%) | 0 / 19 (0%) | 8 / 29 (28%) | 7 / 17 (41%) |
| SAD (%) | 0 / 20 (0%) | 0 / 20 (0%) | 0 / 19 (0%) | 2 / 29 (6.9%) | 1 / 17 (5.9%) |
| Trichotillomania (%) | 0 / 20 (0%) | 0 / 20 (0%) | 0 / 19 (0%) | 1 / 29 (3.4%) | 0 / 17 (0%) |
| Medication | | | | | |
| SSRI (%) | 0 / 20 (0%) | 0 / 20 (0%) | 0 / 19 (0%) | 8 / 29 (28%) | 9 / 17 (53%) |
| Anxiolytic (%) | 0 / 20 (0%) | 0 / 20 (0%) | 0 / 19 (0%) | 2 / 29 (6.9%) | 2 / 17 (12%) |

[1] Mean ± SD, (range Range); n / N; Mean ± SD; n / N (%)

*In-lab MEG sample factor analysis*

Participants completed seven psychiatric questionnaires targeting constructs which are associated with OCD and related disorders: impulsivity, obsessive-compulsivity, perfectionism, intolerance of uncertainty, anxiety, and depression (see Supplementary Fig.

2A). A three-factor solution was found to best explain the latent structure of the item-level questionnaire responses (see Supplementary Fig. 2B). Based on the structure of the individual item loadings (see Supplementary Fig. 2C), we interpret these as an anxious-depressive (AD) factor, an obsessive-compulsive (OC) factor and an intolerant of uncertainty- perfectionistic (IUP) factor. The highest loadings for the AD factor were from the BDI and STAI questionnaires. In the case of the OC factor, OCI-R and PIWSUR predominated. IUS and FMPS items were highest for the remaining IUP factor.

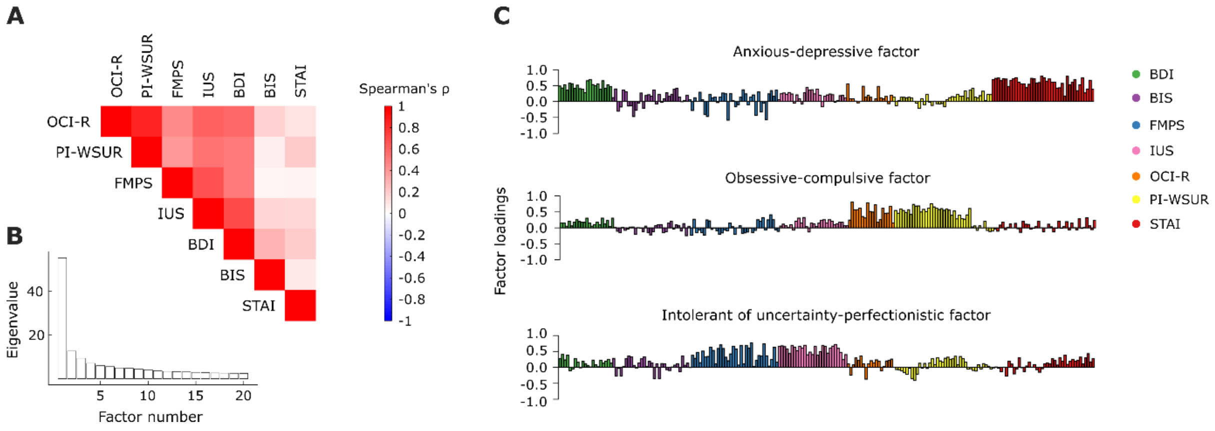


*Supplementary Figure 2: Three latent factors account for the structure of the questionnaire item variance. **A** Correlation matrix of the seven psychiatric questionnaires on which the EFA was based: the Obsessive-Compulsive Inventory – Revised (OCI-R), the revised Padua Inventory questionnaire (PI-WSUR), the Frost Multidimensional Perfectionism Scale (FMPS), the Intolerance of Uncertainty Scale (IUS), the Beck Depression Inventory II (BDI), the Barratt Impulsiveness Scale (BIS), and the State and Trait Anxiety Inventory (STAI). **B** The scree plot for the factor analysis suggests a three-factor solution is optimal. **C** Item loadings onto each of the three factors, an anxious-depressive factor, an obsessive-compulsive factor and an intolerant of uncertainty- perfectionistic factor, colour-coded by the corresponding questionnaire.*

#### *In-lab OCD MEG sample individual differences analysis*

In order to assess whether attenuated ΔES weights were also linked to symptom severity, we correlated the individual beta weights with the scores from the clinical Y-BOCS interview (only conducted in OCD patients), as well as the OCI-R subscale scores, which were available for all in-lab participants as well as for the smartphone population sample. In OCD patients, the effect appears most tightly linked to obsessions, however, this finding is less specific in non-OCD patients across both samples (see Supplementary Fig. 3A-B).

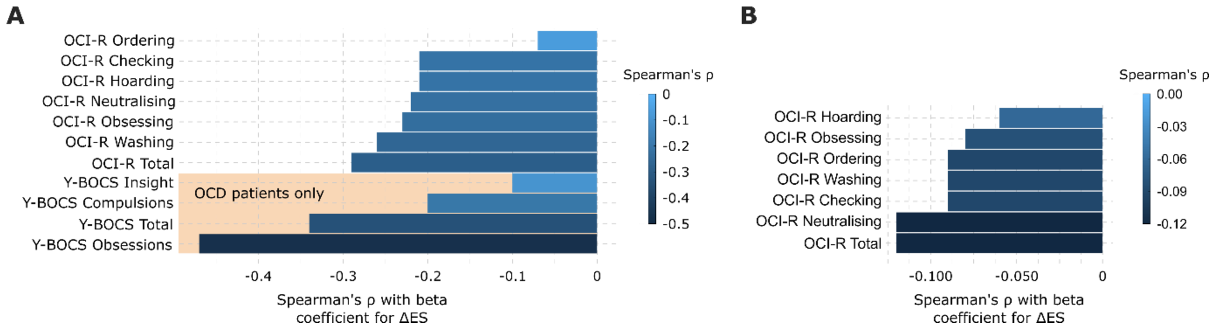


*Supplementary Figure 3: Correlation of the beta weight for ΔES derived from GLMs fit to individual participants and the OCI-R subscales in* ***A*** *the in-lab MEG sample (N = 92 after outlier exclusion based on the GLM beta weights) and* ***B*** *the smartphone population sample (N = 3,903 after outlier exclusion based on the GLM beta weights). Subfigure* ***A*** *additionally comprises the correlations between the GLM beta weight for ΔES and the subscales for the Y-BOCS in the subset of OCD patients, the only group which completed this additional assessment (N = 27 after outlier exclusion based on the GLM beta weights).*

#### *Optimisation of the decoding pipeline on an orthogonal regressor*

We validated the decoding approach and fine-tuned the regularization hyperparameter λ for all subsequent analyses using 5-fold cross-validation (see Supplementary Fig. 4A) on a variable-of-no-interest, namely the current evidence for yellow relative to blue gems at draw d (correlation between current evidence for yellow relative to blue gems and ΔES: $r = 2.755 \times 10^{-4}$, $p = 0.920$). We were able to decode the present stimulus (see Supplementary Fig. 4B and C) and determined the optimal value of the regularization hyperparameter λ to be 0.01 using a non-iterative lasso regression based on all available sensors. However, note that the decodability remains extremely similar for a range of λ values using the iterative lasso regression approach (see Supplementary Fig. 4D).

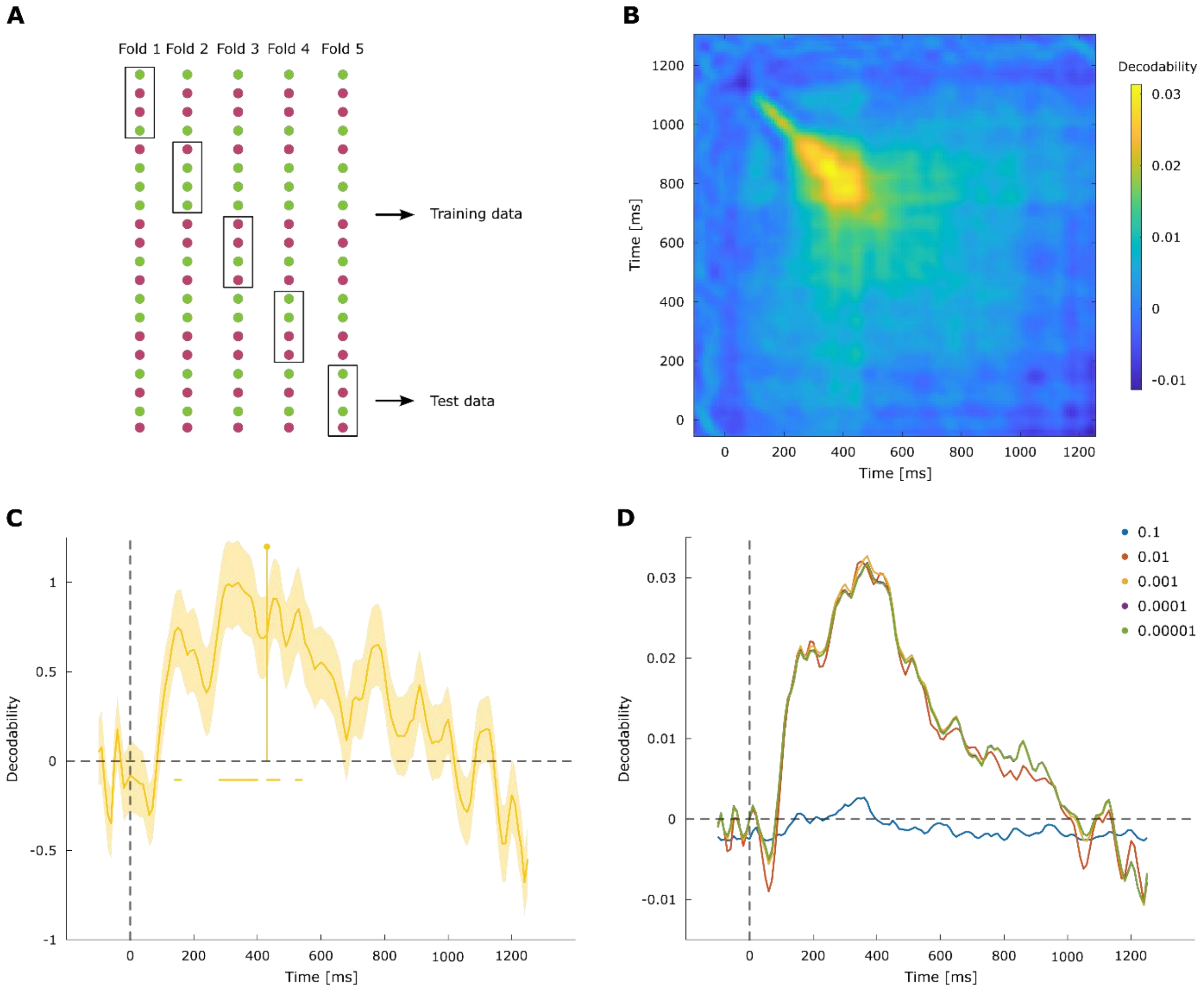


*Supplementary Figure 4: **A** Schematic of the 5-fold cross-validation procedure used to quantify the decodability. Decodability of the current evidence for yellow relative to blue gems over time, quantified as the correlation between trained and tested datapoints using the 5-fold cross-validation procedure for the iterative lasso regression, and visualised in **B** as the complete correlation matrix and **C** as the timecourse of the diagonal. Correlation values were normalized by dividing these by the maximum correlation value for visualization purposes. **D** Decodability of the current evidence for yellow relative to blue gems with the iterative lasso regression approach remains extremely similar across a range of regularisation parameter values.*

#### *Neural ΔES representation*

Following up on the finding of an attenuated neural ΔES representation in high OC participants, we conducted two additional analyses to evaluate whether this association is unique or could be explained by other factors. First, we tested the effect of the commitment to a final decision on the ΔES results, by conducting the same lasso regression and subsequent GLM predicting the decodability from the three factor scores while omitting the data for the evidence sample immediately preceding the decision. All results are highly consistent, both regarding the decodability curve and the individual differences, such that the OC factor predicts attenuated ΔES representation between 420 and 460 ms and between 530 and 560 ms, as opposed to between 420 and 470 ms and between 530 and 560 ms (see Supplementary Fig. 5A).

Second, one may wonder whether the association between ΔES decodability and OC factor score is dependent on this particular update measure. We again ran an equivalent analysis conducting the same lasso regression and subsequent GLM predicting the decodability from the three factor scores, but this time focusing on the cumulative evidence strength $\Sigma ES_d$ instead of ΔES, and find very similar results with respect to the attenuation of the decodability with high OC factor scores – specifically, the OC factor predicts attenuated $\Sigma ES_d$ decodability between 460 and 490 ms and between 530 and 560 ms (see Supplementary Fig. 5B). Note that $\Sigma ES_d$ is calculated in reference to the current majority and therefore still dependent on the current evidence sample.

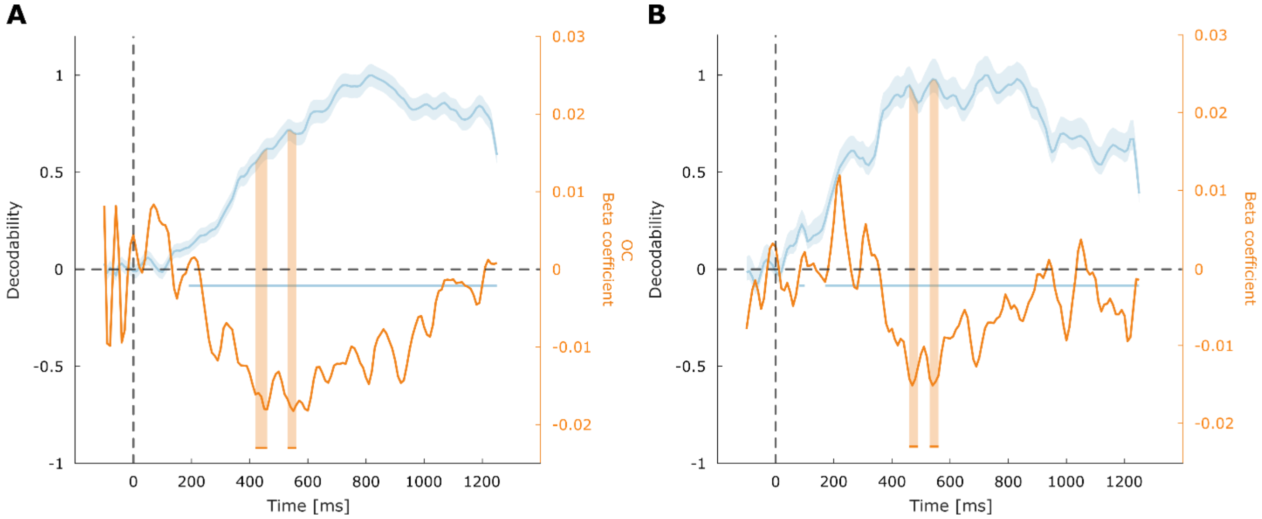


*Supplementary Figure 5: **A** ΔES can be decoded from MEG activity using an iterative multivariate lasso regression also after omitting the data immediately preceding the response (left y axis). Individual differences in OC factor predict significantly decreased decodability within 420-460 ms and 530-560 ms post stimulus (right y axis). **B** Cumulative evidence strength for the current majority $\Sigma ES_d$ can be decoded from MEG activity using an iterative multivariate lasso regression (left y axis). Individual differences in OC factor predict significantly decreased decodability within 460-490 ms and 530-560 ms post stimulus (right y axis).*

*Attenuated neural ΔES representation in the second time window identified through cluster-based permutation testing (530-560 ms)*

The finding of attenuated ΔES decodability is similar within the second time window identified through cluster-based permutation testing of 530-560 ms (see Supplementary Fig. 6A and B). This is also the case for the localisation using searchlight analysis, where the ΔES decodability is widespread, yet the OC effects are primarily evident in mediofrontal regions (see Supplementary Fig. 6C).

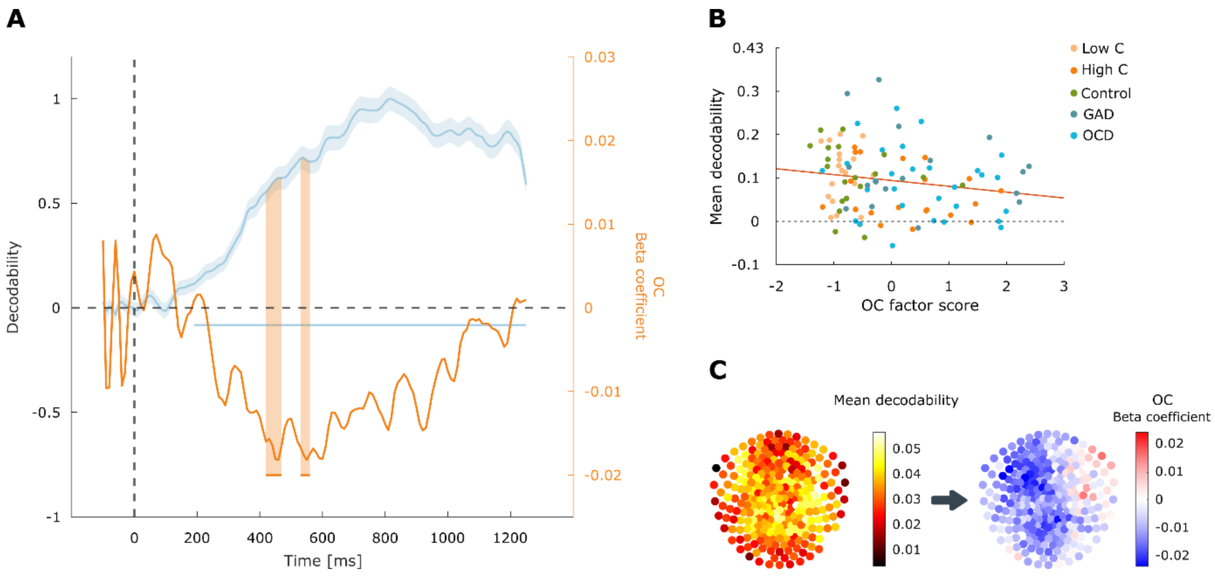


*Supplementary Figure 6: Reduced decodability of ΔES from MEG activity linked to OC symptoms.* ***A*** *ΔES can be decoded from MEG activity using an iterative multivariate lasso regression (left y axis). Individual differences in OC factor predict significantly decreased decodability within 420-470 ms and 530-560 ms post stimulus (right y axis).* ***B*** *Illustration of these attenuated ΔES effects within 530-560 ms.* ***C*** *To investigate the spatial distribution of the sensors contributing to the decodability of ΔES within the time windows where we find individual differences (530-560 ms post stimulus onset), we used searchlight analyses. ΔES are represented in a wider network comprising frontal, mediocentral and occipital areas (column 1). Attenuated ΔES representations in high OC participants were primarily driven by mediofrontal sensors, supporting the idea of altered ΔES processing in mediofrontal areas in high OC participants (column 2).*

#### *Alternative analyses of the localisation of the ΔES representation*

We also investigated the localisation of the ΔES representation using an alternative approach. We calculated the sensor contribution derived from the iterative lasso regression method reported previously[1], whereby we fit the lasso regression fit 2,000 times based on random subsets of 50 sensors instead of the full set of 273 sensors. The contribution of a given sensor was then calculated as the median correlation value for all regression iterations based on sensor subsets including that sensor. To identify any regions which may be particularly associated with reduced decodability of the ΔES, we further fit a GLM predicting the sensor contribution averaged across the time window of interest per sensor from the three factor scores OC symptoms. However, we and others[2] have found this method to yield extremely diffuse maps (see Supplementary Fig. 7), and therefore pursued the searchlight analysis presented in the main text.

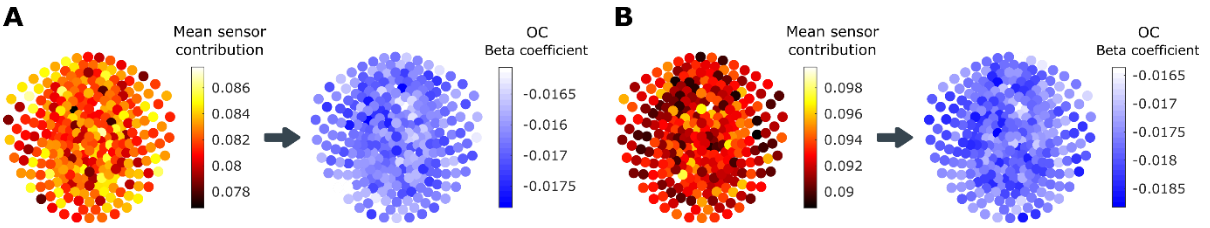


*Supplementary Figure 7: We investigated the spatial distribution of the sensors contributing to the decodability of ΔES within the time windows where we find individual differences (420-470 ms in **A** and 530-560 ms post stimulus onset in **B**). Following the iterative procedure described previously[1], the lasso regression was fit 2000 times based on random subsets of 50 sensors instead of the full set of sensors. The contribution of a given sensor was then calculated as the median correlation value for all regression iterations based on sensor subsets including that sensor (column 1). OC symptoms were linked to decreases in sensor contribution across the board, whereby stronger decreases are depicted with hotter values, as shown by fitting a GLM predicting the sensor contribution averaged across the time window of interest per sensor from the three factor scores (column 2).*

We additionally explored a variant of the iterative lasso regression which omitted the sensor subsetting step, i.e., where the lasso regression was repeated 2,000 times based on the full set of sensors in every iteration. This yielded very similar results to those reported in the main text, albeit with somewhat different time windows for the significant decreases in decodability predicted by the OC dimension, which here are found 350 to 370 ms and 520 to 550 ms post-stimulus (see Supplementary Fig. 8).

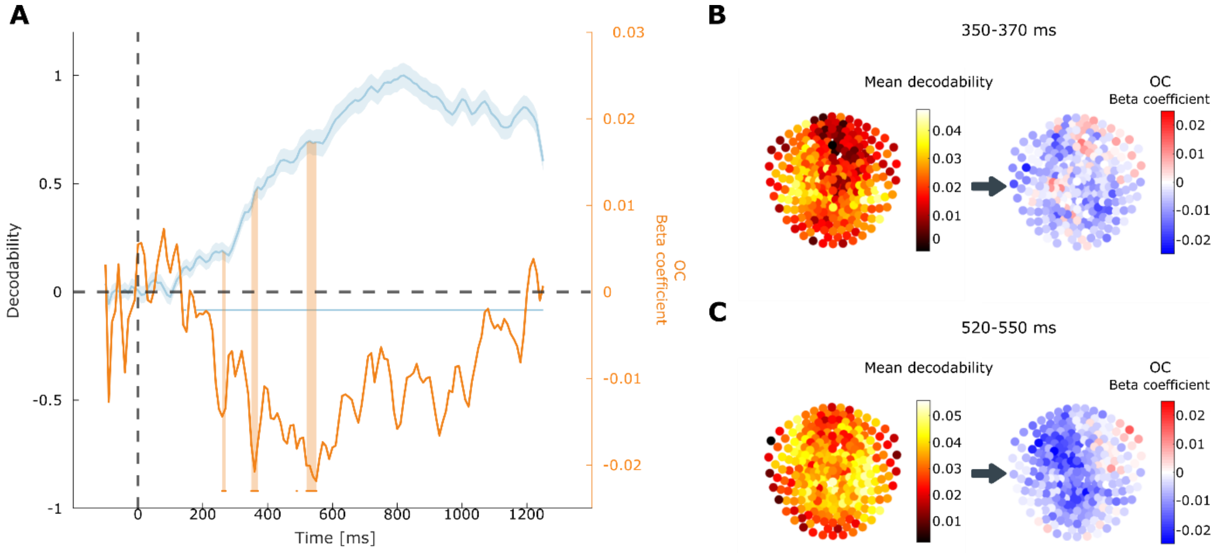


*Supplementary Figure 8: Reduced decodability of the ΔES from MEG activity linked to OC symptoms using a variant of the iterative lasso regression which utilised the full set of sensors on every iteration. **A** ΔES can be decoded from MEG activity (left y axis). Individual differences in OC factor predict significantly decreased decodability within 350-370 ms and 520-550 ms post stimulus (right y axis). **B, C** To investigate the spatial distribution of the sensors contributing to the decodability of ΔES within these shifted time windows where we find individual differences (350-370 ms and 520-550 ms post stimulus), we used searchlight analyses. These largely confirm the findings of ΔES being represented in a wider network comprising frontal, mediocentral and occipital areas (left), and attenuated ΔES representations in OC participants, primarily driven by mediofrontal sensors, particularly in the second more consistent time window (right).*

### References

1. Kurth-Nelson, Z., Barnes, G., Sejdinovic, D., Dolan, R. & Dayan, P. Temporal structure in associative retrieval. *eLife* **4**, e04919 (2015).
2. Rollwage, M. *et al.* Confidence drives a neural confirmation bias. *Nat. Commun.* **11**, 2634 (2020).
